# Spatiotemporally-resolved mapping of extracellular proteomes via *in vivo*-compatible TyroID

**DOI:** 10.1101/2024.08.22.609124

**Authors:** Zijuan Zhang, Yankun Wang, Wenjie Lu, Xiaofei Wang, Hongyang Guo, Xuanzhen Pan, Zeyu Liu, Zhaofa Wu, Wei Qin

## Abstract

Extracellular proteins play pivotal roles in both intracellular signaling and intercellular communications in health and disease. While recent advancements in proximity labeling (PL) methods, such as peroxidase- and photocatalyst-based approaches, have facilitated the resolution of extracellular proteomes, their *in vivo* compatibility remains limited. Here, we report TyroID, an *in vivo*-compatible PL method for unbiased mapping of extracellular proteins with high spatiotemporal resolution. TyroID employs plant- and bacteria-derived tyrosinases to produce reactive *o*-quinone intermediates, enabling the labeling of multiple residues on endogenous proteins with bioorthogonal handles, thereby allowing for their identification via chemical proteomics. We validate TyroID’s specificity by mapping extracellular proteomes and HER2-neighboring proteins using nanobody-directed recombinant tyrosinases. Demonstrating its superiority over other PL methods, TyroID enables *in vivo* mapping of extracellular proteomes, including mapping HER2-proximal proteins in tumor xenografts, quantifying the turnover of plasma proteins and labeling hippocampal-specific proteomes in live mouse brains. TyroID emerges as a potent tool for investigating protein localization and molecular interactions within living organisms.

---

Complex tissues rely on intricate interactions among their various cell types for proper functioning^1^. Extracellular proteins, encompassing both secreted and transmembrane entities, play pivotal roles in orchestrating these interactions, thereby regulating nearly all physiological processes in multicellular organisms through intercellular communication. Consequently, the biochemical characterization of extracellular proteins has yielded significant discoveries, spanning from the identification of peptide hormones to the elucidation of regulators governing physiological functionality^2, 3^. Under pathological conditions, the composition of extracellular proteins can undergo remodeling, exemplified by the dysregulation of ectodomain shedding on neural cell surfaces, which contributes to disease progression^4, 5^. Additionally, a majority of FDA-approved drugs target extracellular proteins^6, 7^, underscoring the importance of developing general methodologies for profiling extracellular proteomes, which would greatly enhance our understanding of cell-cell interactions across various tissues and facilitate the discovery of novel drug targets^8, 9^.

In recent years, there has been remarkable progress in understanding the extracellular proteome landscape, greatly improving our understanding of interactions between cells in various tissues and under different physiological conditions^10, 11, 12^. Notably, proximity labeling (PL) techniques^13^, utilizing Horseradish Peroxidase (HRP), have allowed for precise mapping of extracellular proteomes in model organisms such as flies and mice^14, 15, 16, 17^. This method involves directing HRP to specific cell types, leading to protein labeling through the formation of phenoxyl radicals facilitated by H_2_O_2_. However, due to the significant cytotoxicity of H_2_O_2_, direct administration in living animals is not feasible (**Figure 1a**), necessitating an *ex vivo* approach where the HRP-expressing organ is immersed in a buffered solution containing H_2_O_2_ for protein labeling. Despite the valuable insights gained through this technique, there is a pressing need for non-toxic, *in vivo* methods to label extracellular proteins. Promiscuous biotin ligases, another major type of PL enzyme such as BioID^18^ and TurboID^19^, have been proven effective for intracellular labeling *in vivo*. For instance, promiscuous biotin ligases have been used to map cell type-specific secretomes by labeling in the ER and collecting labeled proteins from a distant region^20, 21, 22, 23^. However, direct *in vivo* labeling of the extracellular proteome faces challenges due to the relatively low concentration of extracellular ATP^24^. Furthermore, both biotin ligases and peroxidase can encounter background interference in cases of high endogenous biotin and H_2_O_2_ levels, respectively^25, 26^. These limitations present significant obstacles to achieving comprehensive profiling of extracellular proteins in living organisms using these approaches.

**Figure 1.**
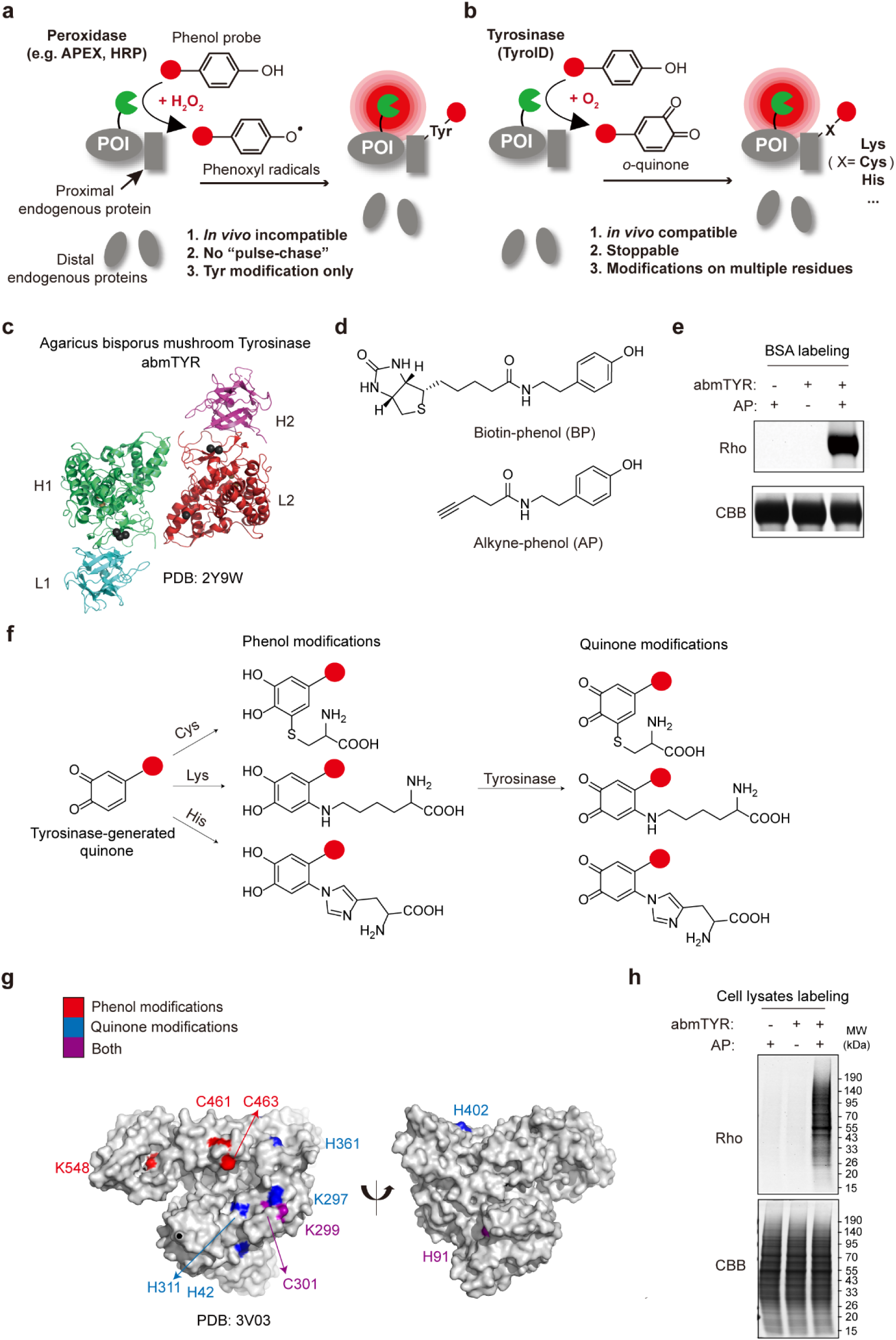
Validation of tyrosinase-based protein labeling *in vitro*. **a.** Schematic of peroxidase-based PL methods such as APEX and HRP. Peroxidase oxidizes phenol probes into reactive phenoxyl radicals with the addition of hydrogen peroxide, which preferentially modifies tyrosines of proximal proteins. The red dot: alkyne or biotin. **b.** Schematic of TyroID. Tyrosinases use oxygen to catalyze the formation of *o*-quinone intermediates from phenol probes, which modifies multiple nucleophilic residues of proximal proteins. The red dot: alkyne or biotin. **c.** The crystal structure of Agaricus bisporus mushroom tyrosinase (abmTYR). **d.** The chemical structures of alkyne-phenol (AP) and biotin-phenol (BP). **e.** The labeling of bovine serum albumin (BSA) by abmTYR and AP *in vitro*. Recombinant BSA was treated with 75 nM abmTYR and 50 μM AP for 30 minutes, followed by click reaction with azide-Rhodamine. The labeling intensity was determined by in-gel fluorescence scanning and Coomassie brilliant blue (CBB) staining was used to demonstrate the equal loading. **f**. Tyrosinase-generated *o*-quinone can react with nucleophilic residues to induce phenol modifications, which can be further oxidized by the tyrosinase to form quinone modifications. **g.** abmTYR-labeled residues on BSA. The residues with phenol and quinone modifications were labeled in red and blue, respectively. Residues with both types of modifications were labeled in purple. **h.** Promiscuous protein labeling by abmTYR and AP in cell lysates. HEK293T cell lysates were treated with 75 nM abmTYR and 50 μM AP for 30 minutes, followed by click reaction with azide-Rhodamine and in-gel fluorescence scanning. Coomassie brilliant blue (CBB) staining was used to demonstrate the equal loading.

Beyond enzymes, the repertoire of small molecule photocatalysts for PL has greatly diversified, now including photosensitizers capable of producing singlet oxygen or phenoxyl radicals upon visible light exposure, enabling cell surface proteome labeling^27^. These photocatalysts comprise a broad range of compounds, such as fluorescent dyes like thiorhodamine^28^ and Eosin Y^29^, as well as co-factors like flavin mononucleotide (FMN)^30^. The Macmillan lab pioneered μMAP, employing an iridium photocatalyst with the unique ability to convert diazirine moieties into highly reactive carbenes via blue light-induced dexter energy transfer^31^. Carbenes, with a lifetime of merely two nanoseconds, result in a more localized labeling compared to phenoxyl radical-based methods, facilitating precise microenvironment mapping. Since these photocatalysts cannot be genetically encoded, they are guided to the target protein on the cell surface through antibodies for PL, greatly aiding in unraveling extracellular proteomes^32, 33^. However, due to the limited penetration of visible light in living organisms, applying these photocatalysts to decipher extracellular proteomes *in vivo* has proven challenging.

In this study, we present TyroID as a novel type of PL technology suitable for non-toxic protein labeling on living cells and in vivo (Figure 1b). TyroID utilizes tyrosinase, a type of enzyme known for catalyzing the oxidation of phenol substrates to o-quinone, a highly reactive electrophile capable of modifying nucleophilic amino acids like cysteines, lysines, and histidines^34,35^. Notably, o-quinone intermediates have previously been utilized in PL applications within both intracellular and extracellular environments, through decaging strategies involving photocatalysts or β-galactosidase^36,37^. However, the ability of tyrosinase to generate quinone species de novo offers a more direct and efficient approach for these biological applications. Unlike other PL methodologies, TyroID only requires the addition of the phenol probe, without the need for supplementary triggers such as H2O2 or visible light. We demonstrate that TyroID can effectively label extracellular proteins without causing toxicity, making it highly promising for in vivo applications. We also validated the spatial specificity of TyroID to map HER2-proximal proteins, discovering Galectin-1 and FUS as novel HER2 neighbors. Finally, we emphasize the versatility of TyroID for in vivo labeling in various organs, including blood, brain, and xenograft tumors.

## Validation of tyrosinase-based protein labeling *in vitro*

To develop TyroID, we began by utilizing Agaricus bisporus mushroom tyrosinase (abmTYR)^38, 39^, a commercially available enzyme commonly used in studies involving the oxidation of tyrosine or dopamine (**Figure 1c**). abmTYR is a tetrameric protein consisting of two H subunits of approximately 392 residues and two L subunits of about 150 residues, with the H subunit exhibiting a fold similar to other tyrosinases, containing a binuclear copper-binding site that catalyzes phenol oxidation^39^. Our initial experiments focused on testing abmTYR’s ability to catalyze the formation of *o*-quinone intermediates for protein labeling using two phenol probes, alkyne-phenol (AP)^40^, and biotin-phenol (BP)^41^, which are commonly used in peroxidase-based PL (**Figure 1d**). Purified bovine serum albumin (BSA) was incubated with abmTYR and AP for 30 minutes. In-gel fluorescence scanning of the labeled BSA, following reaction with azide-rhodamine via Cu(I)-catalyzed azide-alkyne cycloaddition (i.e. click chemistry), revealed significant rhodamine labeling on BSA only in samples treated with both abmTYR and AP (**Figure 1e**). A similar labeling experiment was conducted with BP instead of AP, and streptavidin blotting demonstrated biotinylation on BSA catalyzed by abmTYR (**Figure S1a**).

For a thorough elucidation of the labeled residues on BSA, abmTYR-labeled BSA underwent enzymatic digestion into peptides by Trypsin, followed by analysis using liquid chromatography-tandem mass spectrometry (LC-MS/MS). Tyrosinase-generated *o*-quinone can react with nucleophilic residues to induce phenol modifications, which can then be further oxidized by the tyrosinase to form quinone modifications (**Figure 1f**). Our tandem MS analysis identified both types of modifications on multiple sites, including cysteines, lysines, and histidines, among which several were modified by both types of modifications (**Figure S1b**). The corresponding b and y ions supported the unambiguous assignment of phenol and quinone modifications (**Figure S1c**). Remarkably, these modification sites showed a preference for more surface-exposed residues (**Figure 1g**), possibly due to their higher accessibility, consistent with earlier studies indicating the preference of APEX-generated phenoxyl radicals for surface-exposed tyrosines^42^.

Subsequently, to investigate the versatility of TyroID in labeling various proteins, we treated cell lysates from HEK293T cells with abmTYR and AP, followed by click-based in-gel fluorescence detection. This experimental approach revealed widespread proteome labeling with minimal background in control samples that excluded either the enzyme or AP (**Figure 1h**). The degree of proteome labeling depended significantly on the concentration of each component, showing observable intensity even at concentrations as low as 75 nM for the enzyme and 5 μM for the probe (**Figure S2a, b**). Furthermore, the extent of labeling also varied with the duration of incubation (**Figure S2c**). These findings collectively indicate that tyrosinase-generated *o*-quinone can promiscuously label proteins with high efficiency.

## Evaluation of the live-cell labeling by genetically encoded tyrosinases

We proceeded to establish tyrosinase-based PL in living cells. HEK293T cells were transfected with the catalytic subunit of abmTYR, the H subunit, tagged with a V5 epitope and a nuclear-export sequence (NES). Western blot analysis confirmed the expression of the H subunit in HEK293T cells at the correct molecular weight (**Figure S3a**). However, attempts to initiate intracellular labeling by adding AP to live cells proved unsuccessful, even with supplementation of its cofactor, CuCl_2_ (**Figure S3a**). We speculated that this plant-derived tyrosinase might not function effectively within mammalian cells. Consequently, our attention turned to a bacterial tyrosinase sourced from Bacillus megaterium (BmTYR), which has previously been successfully expressed in mammalian cells^43, 44, 45^. Although the *in vitro* labeling efficiency of BmTYR is much lower than abmTYR (**Figure S3b**), cytosol-resident BmTYR facilitated live-cell proteome labeling with co-treatment of CuCl_2_ and AP (**Figure 2a**). No labeling was observed upon omission of copper, AP or the enzyme, indicating the potential feasibility of genetically encoded tyrosinase for cell type- or organelle-specific PL. Due to the potential toxicity of CuCl_2_, we tested the impact of varying CuCl_2_ concentrations on BmTYR labeling and found that 5 μM is the lowest concentration enabling detectable labeling (**Figure 2a**).

**Figure 2.**
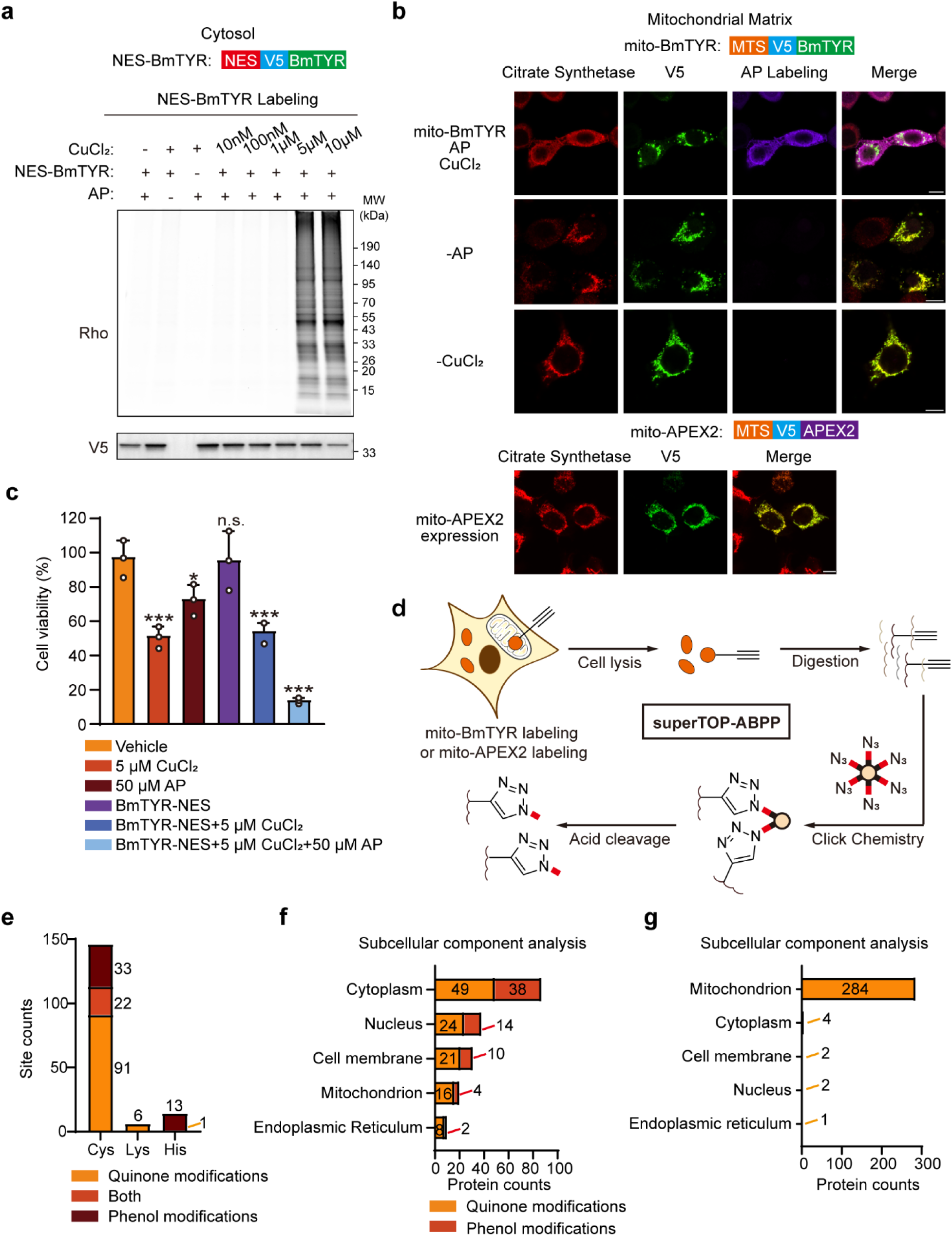
Evaluation of the live-cell labeling by genetically encoded tyrosinases. **a.** Live-cell protein labeling with genetically encoded Bacillus megaterium tyrosinase (BmTYR). HEK293T cells were transfected with NES-BmTYR for 24 hours, treated with CuCl_2_ at various concentrations for 1 hour, followed by the treatment of 50 μM AP for 30 minutes. The cells were lysed and subjected to click-based in-gel fluorescence detection. V5 indicates the expression of BmTYR. **b.** Confocal fluorescence imaging of mito-BmTYR and its impact on mitochondrial morphology. HEK293T cells were transfected with mito-BmTYR for 24 hours, treated with 5 μM CuCl_2_ for 1 hour, followed by the treatment of 50 μM AP for 30 minutes. The cells were fixed and click with azide-biotin after cell fixation to visualize the labeled proteins. Streptavidin-AF647 indicates biotin labeling and Citrate Synthetase indicates the mitochondria. Scale bars, 10 μm. **c.** The impact of CuCl_2_ treatment and BmTYR labeling on cell viability. HEK293T cells were transfected with vehicle or NES-BmTYR for 24 hours, followed by the treatment of 5 μM CuCl_2_ for 1 hour and AP labeling for 30 minutes. The cell viability was determined by the MTS assay. Quantification was performed from three biological replicates and the error bars show mean ±SD. \*\*\**p*<0.001 (student t-test). **d.** The schematic of superTOP-ABPP to identify modification sites by mito-BmTYR and mito-APEX2 labeling. HEK293T cells were transfected with mito-BmTYR or mito-APEX2 for 24 hours and subjected to the corresponding labeling. After digesting the cell lysates into peptides, the labeled peptides were enriched by click reaction with azide-functionalized acid-cleavable agarose beads. The enriched peptides were released by acid cleavage and identified by LC-MS/MS analysis. **e.** The number of cysteines, lysines and histidines labeled by mito-BmTYR with phenol or quinone type of modifications. **f, g**. Subcellular compartment analysis of proteins labeled by mito-BmTYR (**f**) or mito-APEX2 (**g**) with AP. The subcellular information is based on Uniport annotations.

To directly observe the effects of CuCl_2_ on BmTYR and cellular physiology, HEK293T cells were transfected with BmTYR targeted to specific organelles such as the mitochondrial matrix, the ER lumen, and the plasma membrane using organelle-targeting sequences, followed by confocal microscopy imaging of BmTYR and organelle morphology. Unfortunately, we observed aggregation and mis-localization of BmTYR in all compartments, with labeling also lacking adequate subcellular specificity (**Figure 2b and S3c, d**). Furthermore, we noted that Cu supplementation disrupted the morphology of organelle markers, such as Citrate Synthase (CS) for mitochondria, Calnexin for the ER, and Solute carrier family 2, facilitated glucose transporter member 1 (SLC2A1 or GLUT1) for the plasma membrane (**Figure 2b and S3c, d**). Subsequently, a cell viability assay was conducted to quantify the toxicity of Cu addition and BmTYR labeling, revealing that even 5 μM of CuCl_2_ significantly impaired cell viability (**Figure 2c**). Moreover, nearly 90% of cells were dead after BmTYR labeling by treatment with 50 μM AP for 30 minutes. These results collectively suggest that copper supplementation and BmTYR labeling severely disrupt cellular physiology. However, this contrasts somewhat with a recent independent study indicating that BmTYR isn’t toxic during intracellular labeling even under 20 μM CuCl_2_ treatment^46^, a condition that could induce copper-dependent oxidative stress^47, 48^.

We further employed a site-specific chemoproteomic platform, superTOP-ABPP^49^, to map the modification sites of BmTYR labeling in the mitochondrial matrix, which serves as a “gold standard” compartment for assessing the spatial specificity of PL methods^41, 50, 51^ (**Figure 2d**). This approach involved utilizing agarose beads functionalized with azide groups and acid-cleavable linkers to enrich AP-labeled peptides through a single-step click reaction. The enriched peptides, released via acid cleavage, were identified through LC-MS/MS analysis providing site-specific information. In a comparative analysis, we also conducted APEX2 labeling with AP in the mitochondrial matrix and identified its modification sites using the same superTOP-ABPP protocol. The superTOP-ABPP workflow enabled the identification of 146 cysteines, 6 lysines, and 14 histidines modified by AP-derived modifications including both phenol and quinone forms (**Figure 2e and Table S1**). However, subcellular analysis of these labeled proteins revealed an overrepresentation of nuclear and cytosolic proteins with low mitochondrial specificity (**Figure 2f**). In contrast, superTOP-ABPP identified 293 tyrosines labeled by mito-APEX2, and these labeled proteins exhibited high enrichment in the mitochondria (**Figure 2g**). In conclusion, we propose that BmTYR may not be suitable for genetic expression and intracellular protein labeling due to the supplementation of toxic copper.

## Establishment of TyroID for extracellular proteome labeling using recombinant tyrosinases

We therefore aimed to utilize recombinant tyrosinase for extracellular proteome labeling in cultured cells, as Cu ions are naturally bound to the catalytic center during protein purification. MDA-MB-231 and HEK293T cells were treated with 75 nM concentrations of abmTYR along with 50 μM of BP for 30 minutes. Subsequent fluorescent imaging revealed that abmTYR labeling is specific to cell surface and highly co-localizes with the cell surface marker GLUT1 in both cell types, without affecting the localization of GLUT1 (Figure 3a and S4a). We previously found that AP has much better membrane permeability than BP to allow more efficient intracellular labeling^40^. To avoid the potential for AP-derived o-quinone intermediates to penetrate the plasma membrane and label intracellular components, we designed and synthesized a membrane-impermeable alkyne-phenol probe, termed “AxxP”, featuring four polyethylene glycol linkers between the phenol and alkyne groups (Figure 3b and S4b). Fluorescent imaging revealed that abmTYR labeling with AxxP labeling also highly co-localized with GLUT1 in both MDA-MB-231 and HEK293T cells (Figure S4c, d). Further validation through in-gel fluorescence scanning confirmed the promiscuous AxxP labeling of cell-surface proteins, dependent on both the enzyme and the probe (Figure 3c). In comparison, abmTYR generated much higher labeling with AP, likely because AP-derived o-quinone can diffuse into the cells and induce more pronounced labeling on intracellular proteins. Moreover, we also tested AxxP labeling with cell-surface targeted HRP fused with the transmembrane domain of PDGFR (HRP-TM), and significant labeling was observed in an H2O2-dependent manner (Figure S4e).

In contrast to the genetic expression of BmTYR, incubation with recombinant abmTYR and AxxP shows minimal impact on cell viability, even under very high concentrations (**Figure 3d and S4f**). In a direct comparison, MDA-MB-231 cells were exposed to recombinant HRP, and AxxP labeling was triggered by a brief exposure to H_2_O_2_. This resulted in significant cell toxicity in an H_2_O_2_-dependent manner (**Figure 3e**). The non-toxic nature of recombinant tyrosinase for cell surface proteome labeling holds promise for *in vivo* studies and “pulse-chase” experiments that are impractical with HRP labeling. Consequently, we conducted abmTYR labeling on MDA-MB-231 cells for 30 minutes, followed by a 2.5-hour incubation in vehicle medium. The labeling intensity was comparable to that of the sample collected immediately after the 30-minute labeling (**Figure 3f**). Additionally, we performed TyroID labeling in medium containing the enzyme and AxxP for a total of three hours, which resulted in significantly higher labeling intensity. This highlights TyroID’s stoppable nature and its applicability for analyzing the dynamics of labeled proteomes.

**Figure 3.**
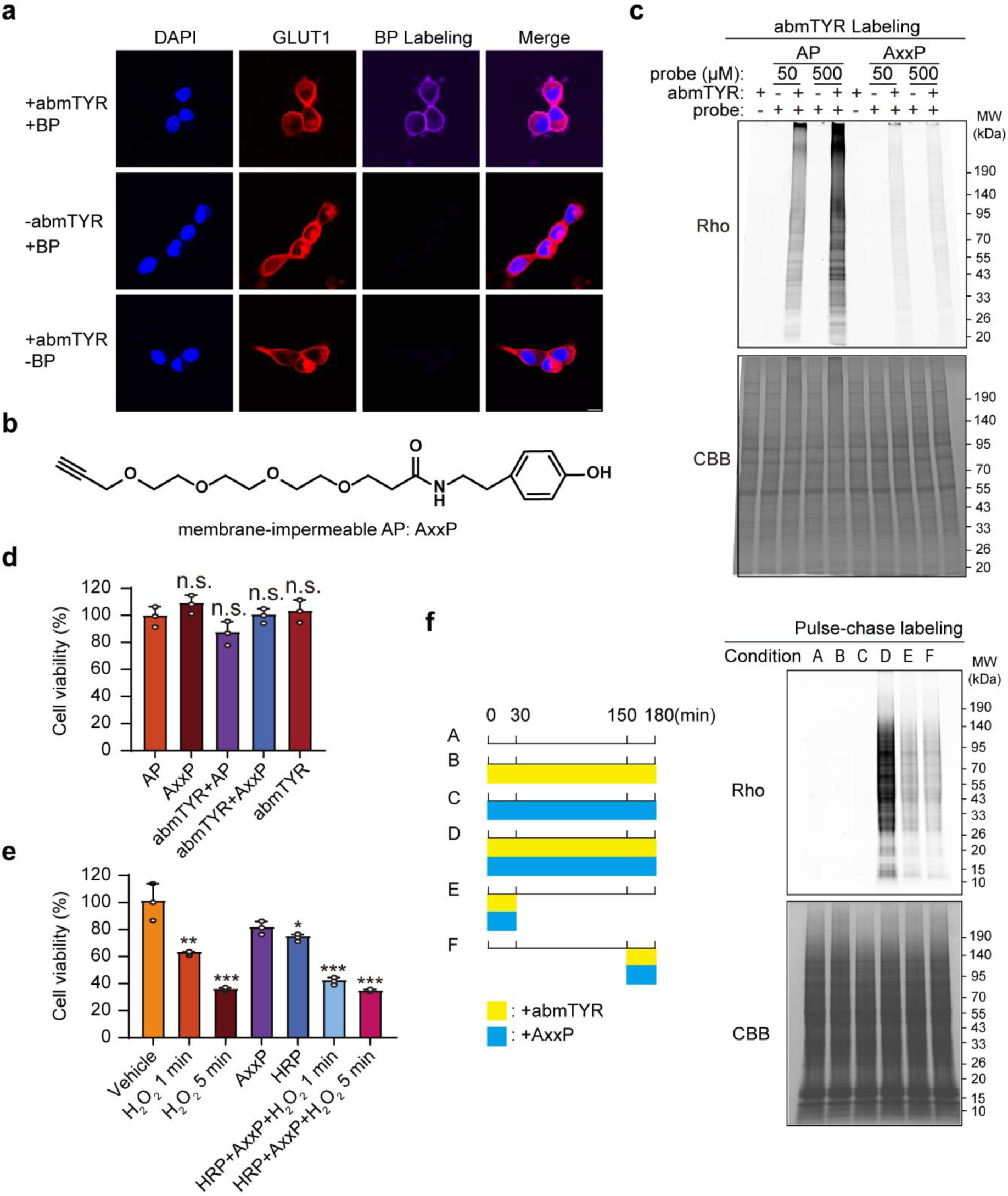
Establishment of TyroID for extracellular proteome labeling using recombinant tyrosinases. **a.** Confocal fluorescence imaging of cell surface proteins labeled by recombinant abmTYR with BP in MDA-MB-231 cells. Streptavidin-AF647 indicates biotin labeling, GLUT1 and DAPI represents the plasma membrane and nucleus respectively. Scale bars, 10 μm. **b.** The chemical structure of membrane-impermeable alkyne-phenol probe (AxxP). **c.** In-gel fluorescence scanning of labeled extracellular proteins by recombinant abmTYR with AxxP and AP. MDA-MB-231 cells were treated with 75 nM abmTYR and 50 or 500 μM of phenol probes (AP and AxxP) for 30 minutes. The cells were lysed and subjected to click-based in-gel fluorescence detection. **d.** The impact of abmTYR-based extracellular labeling on cell viability. MDA-MB-231 cells were treated with 75 nM abmTYR and 50 μM of phenol probes (AP and AxxP) for 30 minutes. The cell viability was determined by the MTS assay. Quantification was performed from three biological replicates and the error bars show mean ±SD. N.S., not significant (student t-test). **e.** The impact of HRP-based extracellular labeling on cell viability. MDA-MB-231 cells were treated with 75 nM HRP and 50 μM AxxP for 30 minutes, followed by 1 or 5 minutes of H_2_O_2_ treatment. The cell viability was determined by the MTS assay. Quantification was performed from three biological replicates and the error bars show mean ±SD. \*\*\**p*<0.001 (student t-test). **f.** Pulse-chase labeling with abmTYR and AxxP. MDA-MB-231 cells were treated with 75 nM abmTYR and 50 μM AxxP for 30 minutes. After washing out the medium, the cells were cultured in vehicle medium for 2.5 hours. The cells were then lysed and subjected to click-based in-gel fluorescence detection. The labeling schematic for all samples is shown on the left.

## Proteomic profiling of extracellular proteins by TyroID

We then conducted quantitative proteomics to identify extracellular proteins labeled by TyroID in MDA-MB-231 cells. The cells were treated with 75 nM of abmTYR and 50 μM of AxxP for 30 minutes, with negative controls omitting either the enzyme or the probe (**Figure 4a**). Following cell lysis, click reaction with azide-biotin was performed, and labeling was confirmed by streptavidin blotting (**Figure S5a**). The biotinylated proteins were subsequently captured by streptavidin beads, followed by on-beads trypsin digestion to release peptides from enriched proteins. These released peptides from each sample were labeled with three different isotopes (“light”, “medium” and “heavy”) using dimethyl labeling before pooling and analysis by LC-MS/MS. The isotopic labels have been swapped across three biological replicates. Peptides were identified, and their dimethyl labeling ratios were quantified using Maxquant. Proteins with at least two peptides were subjected to protein quantification, with the median ratio across all peptides assigned to a specific protein.

**Figure 4.**
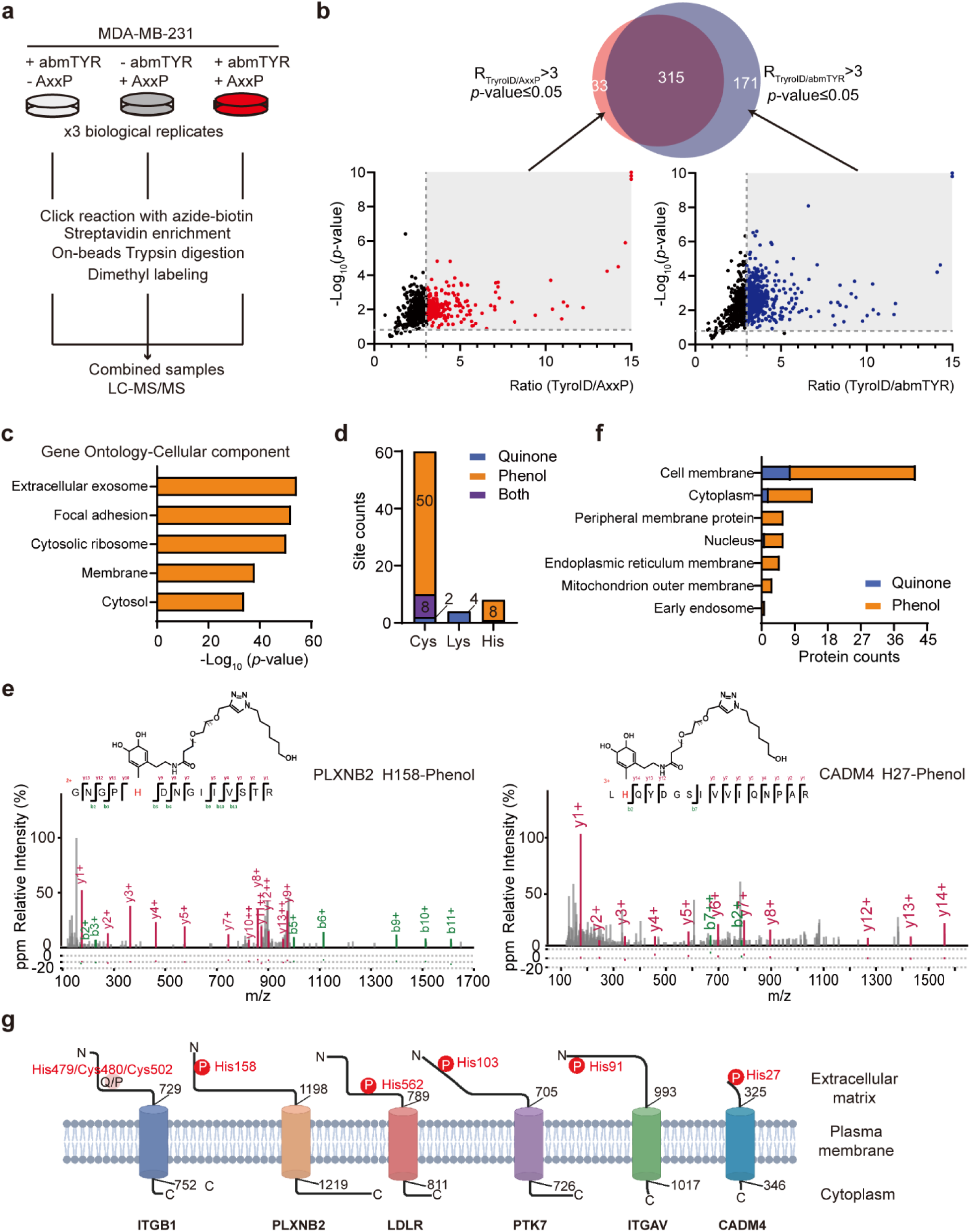
Proteomic profiling of extracellular proteins by TyroID. **a.** The workflow of dimethyl labeling-based quantitative proteomics for mapping extracellular proteins by TyroID. MDA-MB-231 cells were treated with 75 nM of abmTYR and 50 μM of AxxP for 30 minutes, with negative controls that omitted either the enzyme or the probe. The cell lysates were click labeled with azide-biotin, followed by streptavidin enrichment, on-beads trypsin digestion, and dimethyl labeling with three different isotopes. The labeled peptides were mixed and subjected to LC-MS/MS analysis. **b.** Filtering steps for assigning TyroID-labeled proteins. The labeled sample was compared against negative controls omitting either the enzyme or the probe. Proteins with enrichment ratios above 3 and *p-*values below 0.05 in both comparisons were designated as enriched proteins. **c**. GO cellular component analysis of TyroID-enriched proteins. **d.** The number of cysteines, lysines and histidines labeled by abmTYR labeling with AxxP. **e.** Tandem MS spectra of abmTYR-labeled peptides from PLXNB2 and CADM4, with phenol modifications. **f**. Subcellular analysis of site-resolved TyroID-labeled proteins. **g**. TyroID-labeled sites locate at the extracellular regions. The modification sites are labeled in red.

To assign extracellular proteins on MDA-MB-231 cells, we compared the TyroID-labeled sample against each negative control to eliminate potential contaminants. A genuine extracellular protein should display a high enrichment ratio because it should be extensively labeled in the experimental group but not in the control group. Conversely, a false positive, such as an endogenously biotinylated protein or a non-specific bead binder, would exhibit a low ratio since it should be captured similarly in both groups. Thus, proteins with enrichment ratios above 3 and *p*-values below 0.05 in both comparisons were selected to define a total of 315 extracellular proteins (**Figure 4b and Table S2**). Gene Ontology analysis of TyroID-identified extracellular proteins revealed enrichment in extracellular-related terms, such as membrane, focal adhesion, and extracellular exosome, indicating the high specificity of TyroID (**Figure 4c**). We also conducted a global proteomic analysis to assess the impact of TyroID labeling on protein abundance (**Figure S5b and Table S3**). Across 1525 proteins quantified in three biological replicates, none showed significant changes in abundance (**Figure S5c, d**), suggesting that extracellular TyroID labeling does not disrupt cellular homeostasis.

We further employed superTOP-ABPP to map the modification sites of AxxP by abmTYR. This experiment enabled the identification of 60 cysteines, 4 lysines, and 8 histidines modified by AxxP-derived modifications (**Figure 4d and Table S4**), indicating a mass shift of 534.26896 and 536.28461 Da corresponding to the phenol and quinone forms of AxxP modifications, respectively (**Figure 4e and S5e**). These labeled peptides correspond to 60 proteins, fewer than those identified using the dimethyl labeling-based approach, possibly due to challenges in site mapping of AxxP-modified peptides. Subcellular analysis of these site-resolved TyroID-labeled proteins revealed significant enrichment of plasma membrane proteins (**Figure 4f**). Among these proteins, we found six proteins have well-characterized membrane topology information including Integrin beta-1 (ITGB1), Plexin-B2 (PLXNB2), Low-density lipoprotein receptor (LDLR), Inactive tyrosine-protein kinase 7 (PTK7), Integrin alpha-V (ITGAV) and Cell adhesion molecule 4 (CADM4). Interestingly, all TyroID-labeled sites were in their extracellular regions (**Figure 4g**), suggesting the potential of TyroID to provide topological information about transmembrane proteins.

## Cell-selective extracellular protein labeling via HER2-targeting TyroID

To selectively display tyrosinase on the surface of a particular cell type, we employed an antibody-targeting strategy toward the human epidermal growth factor receptor 2 (HER2), a well-characterized oncogene target of high therapeutic interest^52, 53^. We constructed and purified a chimeric enzyme, ZHER-BmTYR, in which BmTYR was fused with ZHER, a nanobody specific for HER2^37^, with well-characterized structural information (**Figure S6a**). We also attempted to purify ZHER fused with the H subunit of abmTYR, but it proved insoluble when expressed in *E. coli* cells (**Figure S6b**). Next, we incubated ZHER-BmTYR with wild-type MDA-MB-231 cells (HER2-negative) or MDA-MB-231 cells stably expressing HER2-green fluorescent protein (GFP) (HER2-positive MDA cells), followed by treatment with AxxP for 30 minutes (**Figure 5a**). We observed that ZHER-BmTYR specifically bound to and labeled the surface of HER2-positive cells, rather than HER2-negative cells (**Figure S6c**). The labeled cells were lysed and subjected to click-based streptavidin blotting, revealing much higher promiscuous proteome labeling of surface proteomes in HER2-positive cells (**Figure 5b**). Additionally, we performed flow cytometry analysis of MDA-MB-231 cells labeled by ZHER-BmTYR and BP after staining with streptavidin-Alexa Fluor 647. HER2-positive MDA cells exhibited significant biotin labeling, with the labeling extent increasing gradually with higher concentrations of ZHER-BmTYR (**Figure 5c and Figure S6d**). Conversely, HER2-negative MDA cells showed detectable biotinylation only under high ZHER-BmTYR concentrations (**Figure 5d**), indicating potential non-specific binding of excess ZHER-BmTYR to HER2-negative cells. These results collectively support the high specificity of the antibody-targeting strategy for displaying TyroID on specific cell types.

**Figure 5.**
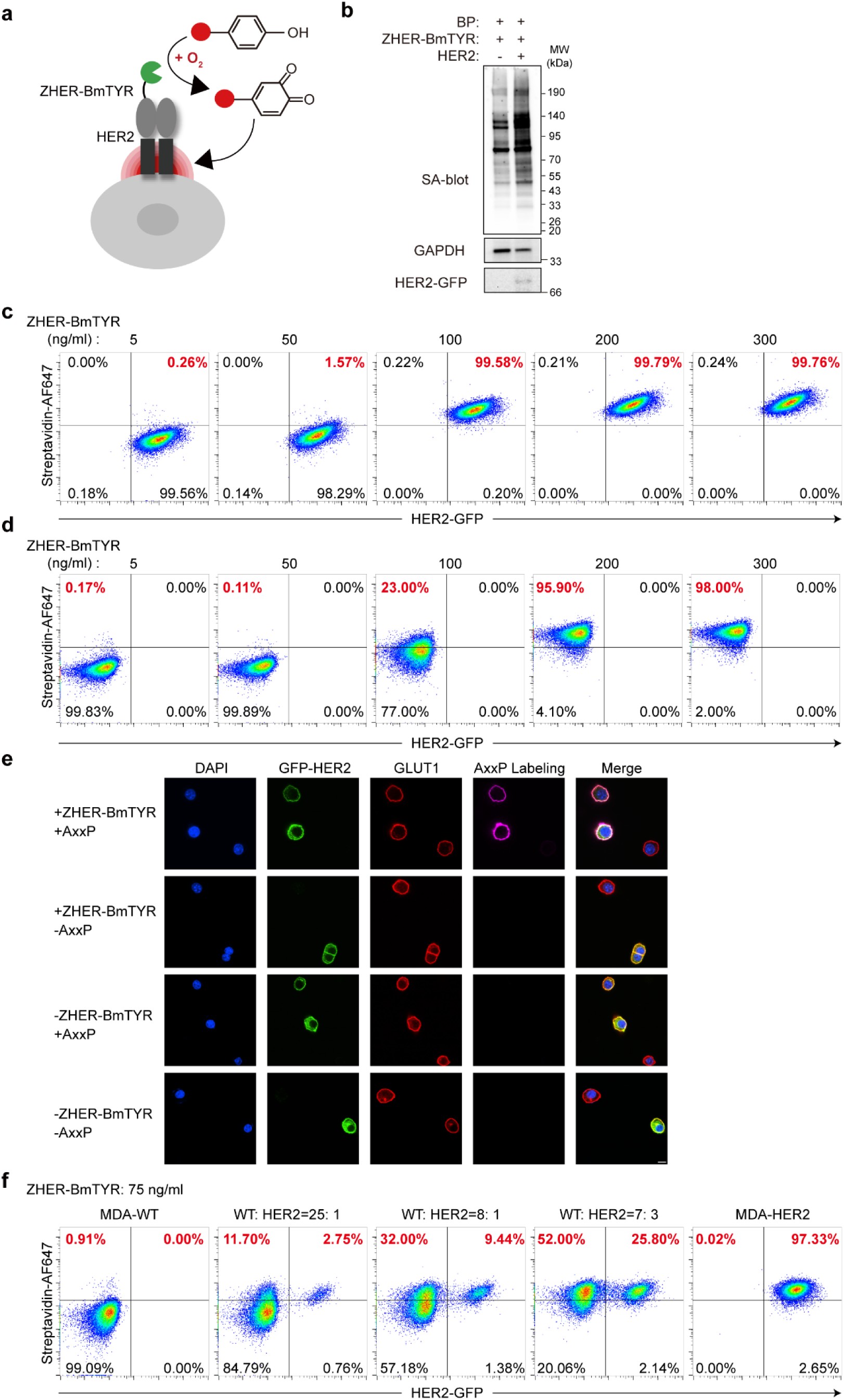
Cell-selective extracellular protein labeling via HER2-targeting TyroID. **a.** The schematic of nanobody-directed cell surface labeling of HER2-expressed MDA cells. **b.** Streptavidin blotting of ZHER-BmTYR labeling on HER2-positive and HER2-negative MDA cells. HER2-positive and HER2-negative MDA cells were treated with 75 nM abmTYR and 50 μM AxxP for 30 minutes. The cells were lysed and subjected to click reaction with azide-biotin, followed by in-gel fluorescence detection of GFP expression and streptavidin blotting of ZHER-BmTYR labeling. **c, d.** Flow cytometry analysis of ZHER-BmTYR labeling on HER2-positive (**c**) and HER2-negative (**d**) MDA cells at different enzyme concentrations. Biotinylation was stained with streptavidin-AF647. **e.** Confocal fluorescence imaging of abmTYR labeling in a co-culture system. HER2-positive and HER2-negative MDA cells were mixed at a 1:20 ratio, followed by the treatment of 75 nM abmTYR and 50 μM AxxP for 30 minutes. The cells were clicked with azide-biotin and fixed to visualize the labeled proteins with streptavidin-AF647. Scale bars, 10 μm. **f.** Flow cytometry analysis of the mixture of HER2-positive MDA cells preincubated with WT MDA cells at various ratios, which was treated with ZHER-BmTYR and BP and stained with streptavidin-AF647. GFP indicates the enzyme expression level and streptavidin indicates the cell labeling level.

We then employed TyroID in a coculture setup by mixing HER2-positive and WT MDA cells. These cocultured cells were incubated with ZHER-BmTYR, treated with BP, and stained with streptavidin-Alexa Fluor 647. Confocal fluorescent imaging reveals the specific biotinylation of HER2-positive MDA cells but not the WT cells (**Figure 5e**). Of note, the distribution of HER2 was not affected by the HER2-targeted TyroID labeling. We further performed the flow cytometry analysis and found that, alongside effectively self-labeling bait cells (HER2-positive), BmTYR-generated *o*-quinone diffused to and labeled neighboring WT cells (**Figure 5f**). The labeling extent of HER2-negative cells increased with a higher ratio of HER2-negative to positive cells, suggesting that labeling of negative cells might be induced by cell-cell contacts. This finding aligns with other PL methods that capture not only the cell surface proteome of bait cells but also reveal cell-cell interactions^54, 55^. Therefore, TyroID holds promise for capturing neighboring cells for subsequent characterizations and deciphering cell-cell interactions in living systems.

## Proteomic profiling of HER2-neighboring proteins by TyroID

Following the successful targeting of BmTYR to HER2 on the cell surface, we further assessed the spatial specificity of TyroID by conducting quantitative proteomics to identify HER2-proximal proteins. HER2-positive MDA cells were incubated with 75 nM of ZHER-BmTYR and 50 μM of AxxP for 30 minutes to initiate PL. We additionally incorporated a negative control that omitted enzyme treatment and a spatial reference, which nonspecifically labeled all extracellular proteins using nanobody-lacking BmTYR (**Figure 6a**). As suggested in prior studies utilizing PL to map interactomes, such a spatial reference is imperative to exclude proteins localized in the same compartment as the bait that may also undergo labeling^56, 57^. After validating the labeling through streptavidin blotting (**Figure S7a**), the samples underwent dimethyl labeling-based quantitative proteomics.

**Figure 6.**
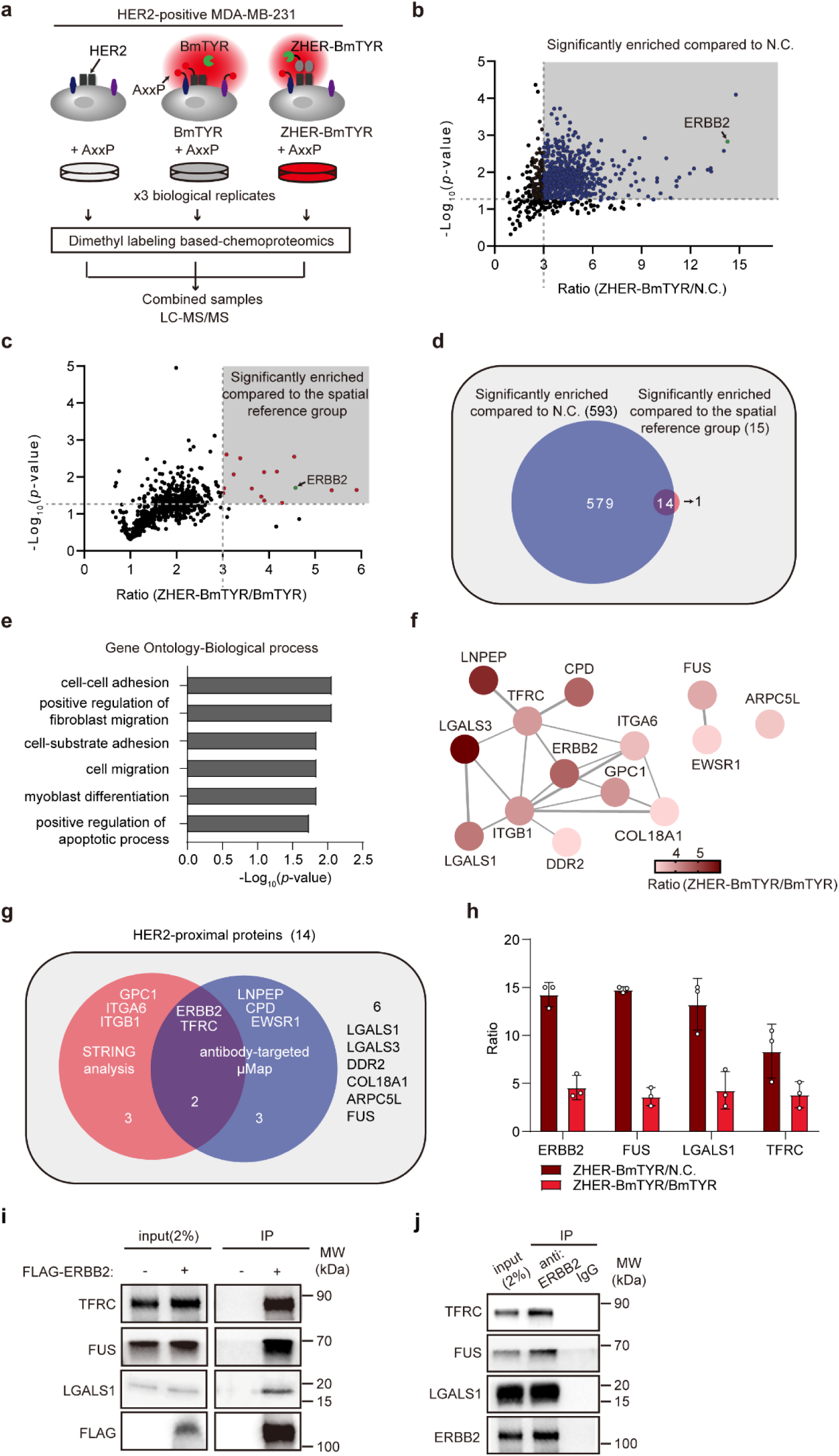
Proteomic profiling of HER2-neighboring proteins by TyroID. **a.** The schematic of nanobody-directed and untargeted TyroID labeling used for mapping HER2-neighboring proteins. HER2-positive MDA cells were treated with 75 nM of ZHER-BmTYR and 50 μM of AxxP for 30 minutes, with a negative control that omitted the enzyme and a spatial reference treated with untargeted BmTYR. The cell lysates were click labeled with azide-biotin, followed by streptavidin enrichment, on-beads trypsin digestion, and dimethyl labeling with three different isotopes. The labeled peptides were mixed and subjected to LC-MS/MS analysis. **b.** Comparison of TyroID-labeled sample against the negative control (N.C.). Proteins with enrichment ratios above 3 and *p*-values below 0.05 are assigned as enriched proteins. **c.** Comparison of TyroID-labeled sample against the spatial reference. Proteins with enrichment ratios above 3 and *p*-values below 0.05 are retained. **d.** Overlap of significantly enriched proteins compared to the negative control and the spatial reference. The overlapping proteins are assigned as HER2-neighboring proteins. **e**. GO biological process analysis of HER2-neighboring proteins. **f**. String analysis of HER2-neighboring proteins revealing their interaction networks. Medium confidence (0.400) was used in the String analysis. **g**. Comparison of HER2-neighboring proteins identified by TyroID and μMAP. **h.** Enrichment ratios of representative HER2-proximal proteins (ERBB2, FUS, LGALS1 and TFRC) compared to the negative control and the spatial reference. Quantification was performed from three biological replicates and the error bars show mean ±SD. **i-j.** Validation of HER2-interacting proteins by co-immunoprecipitation. **i.** MDA-MB-231 cells were transfected with Flag tagged HER2 (i.e. Flag-ERBB2), followed by anti-Flag immunoprecipitation. **j.** Endogenous HER2 in SK-BR-3 cells was immunoprecipitated using protein G beads conjugated with an anti-ERBB2 antibody. The eluates were blotted against FUS and LGALS1, as well as a known HER2-interacting protein TFRC.

We firstly compared the TyroID-labeled sample against the negative control to eliminate potential contaminants, considering proteins with a enrichment ratio above 3 and *p*-values below 0.05 as high-confidence enriched proteins (**Figure 6b**). Gene Ontology analysis of these enriched proteins also revealed terms related to the extracellular environment, such as membrane, focal adhesion, and extracellular exosome (**Figure S7b**). Next, we compared the TyroID-labeled sample against the spatial reference to exclude extracellular proteins distant from HER2. Among the enriched proteins, we retained those with a fold change above 3 and *p*-values below 0.05 (**Figure 6c**), resulting in the identification of fourteen HER2-neighboring proteins (**Figure 6d and Table S5**). As expected, ERBB2 (HER2) is one of the most enriched proteins in both comparisons, suggesting the high spatial specificity (**Figure 6b-c**).

GO analysis of these HER2-neighboring proteins regarding their biological processes exhibits the enrichment of cell-cell adhesion and regulation of fibroblast migration (**Figure 6e**). String analysis reveals a cluster of proteins that are known to have functional links to HER2 (**Figure 6f**), including Integrin alpha-6 (ITGA6), Transferrin receptor protein 1 (TFRC), and Glypican-1 (GPC1). We also compared our dataset with the HER-interacting protein identified by antibody-targeted μMAP^58^ and found five overlapped proteins (**Figure 6g**). The remaining six HER2-neighboring proteins lack direct literature evidence supporting their physical association with HER2. However, some novel hits, such as Galectin-1 (LGALS1), are relevant to HER2 biology or HER2-positive cancer^59^ (**Figure 6h**). For validation, MDA-MB-231 cells were transfected with Flag-tagged HER2, followed by anti-Flag immunoprecipitation and Western Blotting against HER2-proximal proteins. We found that the novel HER2-proximal proteins, LGALS1 and RNA binding protein FUS, interact with HER2 to a similar extent as the known interactor TFRC (**Figure 6i**). Immunoprecipitation of endogenous HER2 further confirmed its interaction with LGALS1 and FUS (**Figure 6j**). Their spatial association with HER2 could unveil novel mechanisms underlying breast cancer progression.

In a parallel comparison, we generated a recombinant APEX2 fused with ZHER, and the *in vitro* labeling assay indicates that the ZHER nanobody does not interfere with APEX2 activity (**Figure S8a, b**). Fluorescence imaging confirms that ZHER-APEX2 specifically binds to and labels the surface of HER2-positive MDA cells (**Figure S8c**). In-gel fluorescence reveals that ZHER-APEX2 generates stronger protein labeling on the cell surface than ZHER-BmTYR (**Figure S8d**), implying that APEX2 produces a greater amount of reactive species. Using the same proteomic workflow and data analysis procedure, ZHER-APEX2 identified 71 HER2-proximal proteins (**Figure S8e**), with ten overlapping proteins identified by ZHER-BmTYR (**Figure S8f**). Only 50.7% of APEX2-identified HER2-proximal proteins are known to interact with HER2, and the remainder might include novel interacting proteins and non-specific bystanders (**Figure S8g**). This proteomic comparison further suggests that APEX2 generates more reactive species and labels more proteins than TyroID. Taken together, these data suggest that TyroID might have a more restricted labeling radius than APEX2 and is highly specific to resolve protein-protein interactions.

## *In vivo* labeling applications enabled by TyroID

The non-toxic and rapid PL facilitated by tyrosinases encouraged us to explore its application *in vivo*, a realm inaccessible to peroxidase-based and visible light-activated PL methods. Therefore, we conducted three distinct *in vivo* experiments to demonstrate its efficacy in living mice. Initially, we sought to analyze HER2-neighboring proteins *in vivo* rather than in cell cultures. To achieve this, we established a xenograft mouse model by injecting HER2-positive MDA cells into nude mice. Subsequently, mice were intratumorally injected with ZHER-BmTYR and AxxP to initiate labeling (**Figure 7a**). After a 12-hour period, tumor tissues were harvested and subjected to click labeling with azide-rhodamine. Notably, robust fluorescence signals were detected in labeled tumors, whereas those treated solely with AxxP exhibited markedly lower fluorescence intensity (**Figure 7b**). Furthermore, tumor lysates underwent click labeling with azide-biotin, revealing extensive proteome biotinylation as evidenced by streptavidin blotting, thus confirming successful TyroID labeling of tumors in living mice (**Figure 7c**). Subsequently, the labeled proteins were enriched and subjected to quantitative proteomics using dimethyl labeling (**Figure S9a**). This *in vivo* experimentation demonstrated a correlated extent of enrichment of HER2-neighboring proteins akin to that observed in cell culture settings, underscoring the high specificity of *in vivo* TyroID labeling (**Figure 7d and Table S6**). Moreover, *in vivo* labeling unveiled several novel HER2-neighboring proteins, previously unrecognized in cell culture-based experiments, such as Transitional endoplasmic reticulum ATPase (VCP), Calponin-2 (CNN2), and Sorting nexin-9 (SNX9) (**Figure 7e**). Interestingly, VCP or p97 is involved in the ER-associated protein degradation (ERAD) process, and a previous study suggested that VCP expression significantly correlates with poor survival in HER2-positive, but not HER2-negative, breast cancer patients^60^. The association between VCP and HER2 during tumorigenesis might provide novel mechanisms underlying its phenotypic consequences.

**Figure 7.**
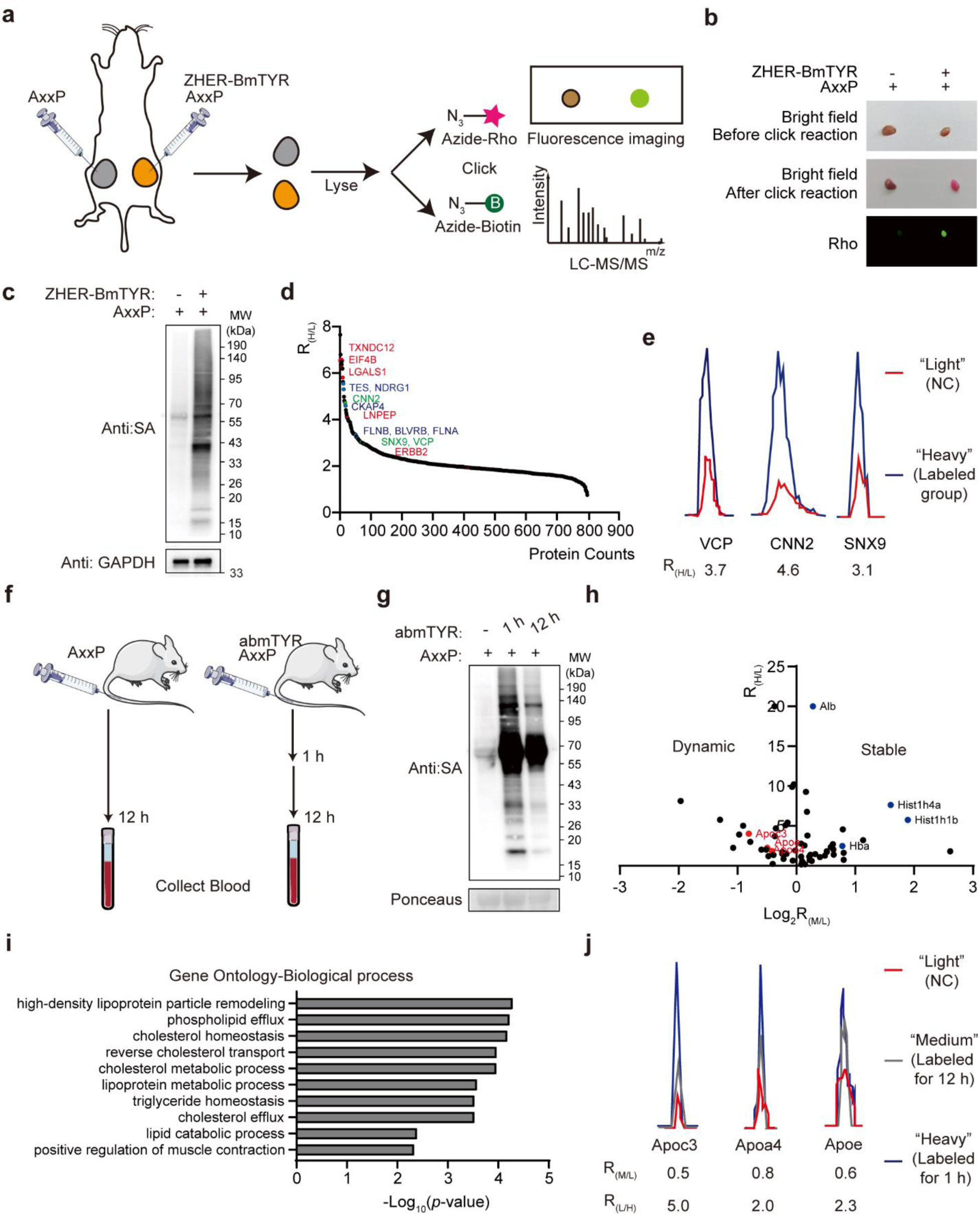
*In vivo* TyroID labeling of HER2-neighboring proteins in tumor xenografts and circulating proteins in the blood. **a.** The schematic of HER2-targeting TyroID labeling in tumor xenografts. Tumor xenografts were established by injecting HER2-positive MDA cells into nude mice, which were intratumorally injected with ZHER-BmTYR and AxxP for 12 hours. **b.** Fluorescence imaging of ZHER-BmTYR labeling in tumor tissues. The labeled tumor tissues were subjected to click reaction with azide-rhodamine and the rhodamine intensity was detected. **c.** Streptavidin blotting of ZHER-BmTYR labeling in tumor xenografts. The labeled tumor tissues were lysed and subjected to click reaction with azide-biotin, followed by streptavidin blotting. **d.** The distribution of protein enrichment ratios in TyroID labeling of Tumor xenografts. Representative HER2-neighboring proteins identified in cell culture are colored in red. Representative HER2-neighboring proteins identified by μMAP are colored in blue. Representative novel hits identified in *in vivo* experiments are colored in green. **e.** The representative MS1 chromatographic peaks of peptides from VCP, CNN2 and SNX9. **f.** The schematic of *in vivo* protein labeling in the blood. The mice were injected with abmTYR and AxxP intravenously, followed by the collection of blood in 1 or 12 hours. **g.** Streptavidin blotting of abmTYR labeling in the blood of living mice. Proteins extracted from the blood were click reacted with azide-biotin, followed by streptavidin blotting. **h.** The turnover of TyroID-labeled plasma proteins. Proteins with light/heavy ratio over 2 were assigned as TyroID-labeled proteins. Proteins with medium/light ratio below 0.5 and over 2 were assigned as dynamic and stable plasma proteins, respectively. **i.** GO biological process analysis of dynamic plasma proteins. **j.** The representative MS1 chromatographic peaks of peptides from Apoc3, Apoa4 and Apoe.

For the second *in vivo* experiment, we administered abmTYR and AxxP intravenously into the bloodstream of living mice (**Figure 7f**). Following TyroID labeling, the mice exhibited normal behavior, and blood samples were collected at 1- and 12-hour intervals post-injection. Plasma proteins were extracted and subjected to click reaction with azide-biotin. Streptavidin blotting unveiled extensive labeling of plasma proteins in an enzyme- and probe-dependent manner (**Figure 7g**). Interestingly, we noted that labeling of plasma proteins decreased in the 12-hour sample, which led us to hypothesize that *in vivo* labeled plasma proteins might undergo degradation or translocation to other organs within the 12-hour timeframe. To quantify the turnover rates of various plasma proteins, we performed quantitative proteomics to compare the labeled proteins between 1- and 12-hour treatments, resulting in the quantification of 44 TyroID-labeled plasma proteins (**Figure S9b and Table S7**). We identified a group of proteins showing a dramatic decrease in labeling at 12 hours, while another group of proteins remained quite stable (**Figure 7h**). These stable plasma proteins include high-abundant proteins such as histones and serum albumin that are constantly present in the blood, whereas the dynamic proteins are highly enriched in cholesterol homeostasis and phospholipid efflux (**Figure 7i**). For instance, TyroID identifies three apolipoproteins with fast turnover rates including Apolipoprotein C-III (Apoc3), Apolipoprotein A-IV (Apoa4) and Apolipoprotein E (Apoe) (**Figure 7j**), and they are structural components of plasma lipoprotein that influence vascular biology^61^. Interestingly, the turnover rate of Apoe has been determined in central nervous system using an *in vivo* stable isotope labeling strategy, revealing its fast clearance kinetics^62^. These plasma proteins with fast turnover rates might mediate transient molecular functions in the blood circulatory system.

For the final *in vivo* experiment, we evaluated TyroID labeling in the intact mouse brain by administering abmTYR and BP into the left hippocampus (**Figure 8a**). As an internal control, we solely administered BP into the right hippocampus. Brain tissue was harvested 1-hour post-injection abmTYR and BP to prepare brain slices, followed by staining with streptavidin-Alexa Fluor 647. The biotinylation was prominently enriched at the left injection site with minimal spread to the surrounding area, whereas the right injection site and untreated controls exhibited low background levels (**Figure 8b**). Upon closer examination, we confirmed that the distribution of biotinylation is largely segregated from the nucleus, supporting the high subcellular specificity (**Figure 8c**). The brain slices were also stained for cleaved caspase 3, a molecular marker of apoptosis, which showed undetectable signal, indicating the low toxicity of TyroID labeling *in vivo* (**Figure S9c**). Subsequently, we dissected the hippocampus for streptavidin blotting analysis, confirming selective protein labeling by TyroID in the left hippocampus, with minimal labeling signal in the right hippocampus (**Figure 8d**). Furthermore, quantitative proteomics analysis was performed to identify labeled proteins in the hippocampus (**Figure 8e and S9d**), revealing numerous brain-specific cell surface markers such as Plexin-A1 (Plxna1), Neurochondrin (Ncdn) and Neuronal membrane glycoprotein M6-a (Gpm6a) (**Figure 8f and Table S8**). GO cellular component analysis also reveal the enrichment of brain-specific terms including myelin sheath, synapse, and neuronal cell body (**Figure 8g**). Taken together, TyroID enables *in vivo* extracellular labeling in various systems.

**Figure 8.**
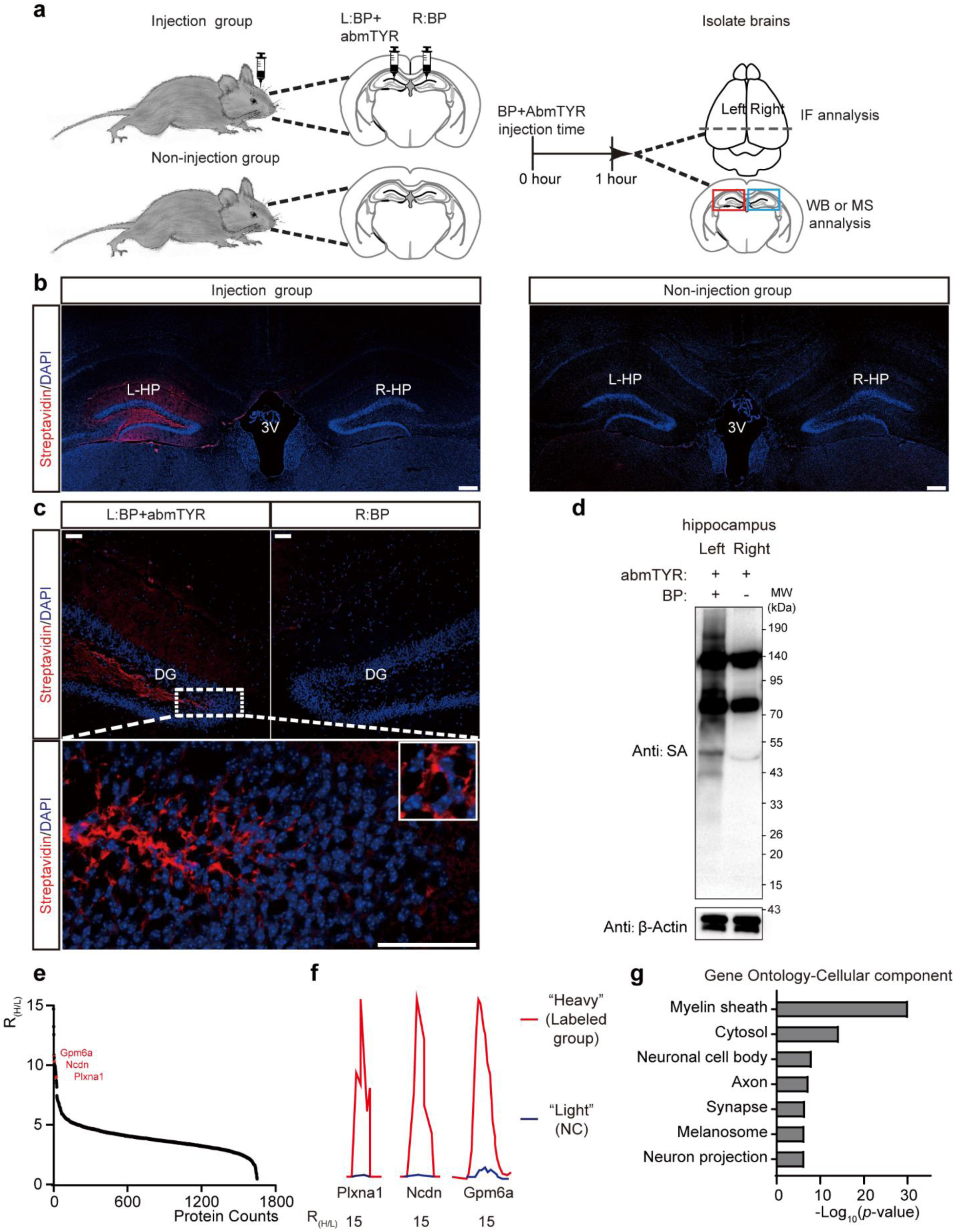
*In vivo* TyroID labeling of extracellular proteins in the hippocampus. **a.** The schematic of *in vivo* TyroID labeling in the hippocampus. Left panel: Illustration of intracranial injection of abmTYR with biotin phenol (BP) into the left (L) hippocampus (HP) and BP alone into the right (R) HP. Right panel: Diagrammatic timeline outlining the experimental protocol, including the injection, follow-up period, and subsequent isolation of brain or hippocampal tissues. (**b-c**) Immunofluorescence staining for streptavidin in the HP (**b**) and dentate gyrus (DG) (**c**) of brains from the injection group and the non-injection group. Nuclei are labeled with DAPI. Insets display magnified views of the regions indicated. 3V, third ventricle. Scale bars: (**b**) 100 μm; (**c**) 50 μm. **d.** Streptavidin blotting of abmTYR labeling in the hippocampus of living mice. The left and right hippocampus were dissected and analyzed by streptavidin blotting. **e.** The distribution of protein enrichment ratios in hippocampus-targeted TyroID labeling. Representative brain-specific cell surface proteins are colored in red. **f.** The representative MS1 chromatographic peaks of peptides from Plxna1, Ncdn and Gpm6a. **g.** GO cellular component analysis of TyroID-labeled proteins in the hippocampus.

## Discussion

PL has become a powerful tool for capturing extracellular protein information in mass spectrometry experiments, especially with the recent use of genetically encoded HRP and antibody-directed photocatalysts to uncover nanoscale arrangements of cell surface proteins^14, 15, 16, 17, 27, 28, 30^. However, until now, profiling extracellular proteomes in living animals has been largely inaccessible with existing PL techniques. The TyroID method presented here offers a fast and non-toxic approach to label endogenous proteins with bioorthogonal handles *in vivo*, thus enabling subsequent chemoproteomic profiling.

TyroID harnesses tyrosinase to catalyze the production of protein-reactive *o*-quinone from phenoxyl probes, akin to substrates employed in peroxidase-based PL. Unlike peroxidase-generated phenoxyl radicals, which predominantly modify tyrosines, *o*-quinone intermediates exhibit high reactivity and can label multiple nucleophilic residues. Recent developments in PL methods targeting amino acids beyond tyrosines and lysines have broadened the scope of labeled proteomes, complementing classical PL approaches^27, 63, 64^. Consequently, TyroID may label proteins lacking exposed tyrosines, evading detection by peroxidases. While quinone derivatives have been utilized for both intracellular and intercellular PL, their labeling radius is expected to exceed that of peroxidase-generated phenoxyl radicals. To enhance the spatial specificity of TyroID, we devised a membrane-impermeable alkyne-phenol probe to prevent the penetration of *o*-quinone intermediates into cells. Beyond cell surface specificity, we have showcased TyroID’s spatial resolution in delineating HER2-proximal proteins through the incorporation of suitable spatial references. However, further endeavors are warranted to engineer modified phenoxyl probes for precise modulation of its diffusion radius.

The primary advantage of TyroID over peroxidase-based and other existing PL methods lies in its compatibility with *in vivo* settings. To illustrate its efficacy in living organisms, we conducted three distinct labeling experiments across various tissue organs, generating proteomic datasets poised to yield valuable biological insights: (1) Leveraging a HER2 nanobody to guide tyrosinase to HER2 on tumor cell surfaces enabled localized PL, resulting in the creation of the inaugural *in vivo* HER2 interactome dataset.

1. (3) Direct labeling of plasma proteins *in vivo*, followed by tracking their turnover. (2) Precise injection of tyrosinase into the brain allowed for hippocampal-specific mapping of extracellular proteomes. Notably, throughout all *in vivo* experiments, mice exhibited no signs of toxicity during the labeling process. Consequently, we posit that TyroID represents an ideal tool for mapping cell type-specific extracellular proteomes or elucidating cell-cell interactions within living organisms.

It’s important to highlight a significant limitation of TyroID compared to traditional PL methods, namely its inability to perform intracellular labeling. While TyroID is genetically encodable and can be targeted to specific subcellular regions post-protein synthesis, its activity necessitates the exogenous addition of copper, which poses toxicity concerns and complicates *in vivo* experimentation. Further safety assessment of genetically encoded TyroID is imperative before its application *in vivo*, such as generating TyroID transgenic mice. One potential solution could involve the delivery of recombinant tyrosinase pre-bound with copper into cells for intracellular labeling using protein delivery strategies^65, 66^.

In a previous study, we developed a proteomic technique termed TransitID, which integrates TurboID and APEX2 labeling to track dynamic proteome trafficking within and between living cells^40^. Presently, this cutting-edge technology is confined to experiments conducted in cell cultures due to challenges associated with utilizing the peroxidase enzyme required for destination labeling *in vivo*. However, given that TyroID also employs non-biotin probe substrates, it presents a viable alternative to APEX2 and operates orthogonally to TurboID labeling. As such, TyroID can seamlessly replace APEX2 in the TransitID framework, enabling the elucidation of proteome trafficking *in vivo* between two cell types that cannot be physically separated by dissection. We anticipate the development of this *in vivo* TransitID technology and its broad applications across various living systems.

## Acknowledgements

This work was supported by the Science Fund Program for Distinguished Young Scholars (Overseas) from National Natural Science Foundation of China, Youth Talent Cultivation Fund of Tsinghua University, Beijing Frontier Research Center for Biological Structure, and the Fundamental Research Funds from Beijing National Laboratory for Molecular Sciences (BNLMS202301). W.Q. is sponsored by Bayer Investigator Award. Z.W. is supported by the Young Elite Scientists Sponsorship Program by Beijing Association for Science and Technology. We thank Dr. Xing Chen at Peking University for kindly providing MDA-MB-231 cells stably expressing GFP-HER2. We thank Dr. Weidi Xiao, Dr. Yuan Liu and Dr. Chu Wang at Peking University for providing guidance on superTOP-ABPP experiments.

## Author contributions

Z.Z., Y.W. and W.Q. designed the research and analyzed all the data except where noted. Z.Z. and Y.W. performed all experiments except where noted. Z.Z., Y.W. and W.Q. designed the proteomics experiments. Z.Z. and Y.W. performed the proteomic samples. Z.Z., Y.W. and X.P. analyzed the MS data. Z.Z. and H.G. performed imaging experiments. W.L. performed protein purification experiments. W.L. and Z.L. performed mouse experiments for the tumor and blood labeling. X.W performed the experiments in mouse brain with the guidance of Z.W.. Z.Z., Y.W. and W.Q. wrote the paper with input from all authors.

## Online methods

### Cell culture

HEK 293T, SK-BR-3 and MDA-MB-231 cells from the ATCC (passages <25) were cultured in DMEM (Thermo Fisher) supplemented with 10% fetal bovine serum (Biological Industries), 100 units/ml penicillin and 100 mg/ml streptomycin at 37 °C under 5% CO_2_. For fluorescence microscopy imaging experiments, cells were grown on 15-mm glass-bottom cell culture dish (NEST). For proteomic experiments, cells were grown on 15-cm glass-bottomed Petri dishes (NEST). For In-gel fluorescence experiments, cells were grown on six-well plates (NEST). MDA-MB-231 cells stably expressing GFP-HER2 are kindly gifted from the Xing Chen lab at Peking University.

### Plasmids

Polymerase chain reactions (PCR) were performed with Phanta DNA Polymerase (catalog no. P520) purchased from Vazyme. Ligase-free cloning reactions were performed with Basic Seamless Cloning and Assembly Kit (catalog no. CU201) purchased from TransGen. Plasmid products were transformed into DH5α competent cell (catalog no. TSC-C14) purchased from Tsingke. DNA sequences were confirmed by Sanger sequencing before use.

The DNA of BmTYR and the catalytic subunit of abmTYR were ordered from Azenta, codon-optimized for human expression. The PCR products with the following sequences were inserted between the EcoRI and AfeI sites of pcDNA3.1 vector to generate NES-abmTYR, NES-BmTYR, Mito-BmTYR and BmTYR-KDEL.

NES-abmTYR: NES-GSGS-V5-GSGS-abmTYR (NES: LQLPPLERLTLD; V5: GKPIPNPLLGLDST) NES-BmTYR: NES-GSGS-V5-GSGS-BmTYR

Mito-BmTYR: MTS-GSGS-V5-GSGS-BmTYR (MTS (COX4): MLATRVFSLVGKRAISTSVCVRAH)

BmTYR-KDEL: Igk-GSGS-V5-GSGS-BmTYR-KDEL (SS (Ig kappa chain): METDTLLLWVLLLWVPGSTGD)

BmTYR-TM was constructed from HRP-TM which was purchased from Addgene (Plasmid #44441) by replacing HRP with BmTYR.

pET28a-ZHER-BmTYR and pET28a-ZHER-abmTYR was constructed from pET28a-ZHER-LacZ (a gift from Xing Chen at Peking University) by replacing LacZ with BmTYR and abmTYR. pET28a-BmTYR was constructed from pET28a-ZHER-LacZ by replacing ZHER-LacZ with BmTYR. pET28a-ZHER-APEX2 was constructed from pET28a-ZHER-LacZ by replacing LacZ with APEX2.

pLX304-Mito-V5-APEX2 was a gift from Chu Wang at Peking University. FLAG-HER2 plasmid (catalog no. P62918) was purchased from Miaolingbio.

### Synthesis of AxxP

AxxP was ordered from Amatek. ^1^HNMR (400 MHz, CDCl_3_) δ 7.03 (m,2H), 6.81 (m, 2H), 6.58 (d, 1H), 4.20 (d, 2H), 3.61 (m.16H), 2.73 (t, 2H), 2.44 (m,3H).

### TyroID labeling *in vitro* and in living cells

For the labeling of abmTYR on BSA, 3 mg/mL BSA was incubated with 75 nM abmTYR and 50 μM AP for 30 minutes. The reaction was stopped by precipitating the BSA. The precipitated BSA was resuspended in 0.4% SDS/PBS and reacted with 100 μM azide-Rhodamine, premixed 2-(4-((bis((1-tertbutyl-1H-1,2,3-triazol-4-yl)methyl)amino)methyl)-1H-1,2,3-triazol1-yl)-acetic acid (BTTAA)-CuSO_4_ complex (500 μM CuSO_4_, BTTAA:CuSO_4_ with a 2:1 molar ratio) and 2.5 mM freshly prepared sodium ascorbate for 2 hours at 25 °C. The excess reagents were removed by methanol and chloroform (4:1) precipitation and the fluorescent intensity was detected by in-gel fluorescence scanning. For the labeling of abmTYR with BP, the labeled BSA was reacted with 100 μM azide-biotin using the same click protocol and analyzed by Western blot.

For the labeling of abmTYR in cell lysates, HEK293T cells were lysed in PBS with sonication and lysates were clarified by centrifugation at 20,000 g for 10 minutes at 4 °C. Protein concentration in clarified lysates was estimated with Pierce BCA Protein Assay Kit (Thermo Fisher) and normalized to 2 mg/mL. Lysates were incubated with abmTYR and AP. 50 μL of lysates were reacted with 100 μM azide-Rhodamine, premixed 2-(4-((bis((1-tertbutyl-1H-1,2,3-triazol-4-yl)methyl)amino)methyl)-1H-1,2,3-triazol1-yl)-acetic acid (BTTAA)-CuSO_4_ complex (500 μM CuSO_4_, BTTAA:CuSO_4_ with a 2:1 molar ratio) and 2.5 mM freshly prepared sodium ascorbate for 2 hours at 25 °C. The excess reagents were removed by methanol and chloroform (4:1) precipitation and the fluorescent intensity was detected by in-gel fluorescence scanning.

For the labeling of genetically-encoded BmTYR in living cells, HEK293T cells were transiently transfected with BmTYR fusion construct of interest for 24 hours. Cells were washed twice with Hank’s Balanced Salt Solution (HBS, catalog no. H1025, Solarbio), and then incubated with 5 μM CuCl_2_ in HBS for 1 hour, followed by the treatment of 50 μM alkyne-phenol for 30 minutes. The reaction was quenched by replacing the HBS with an equal volume of quenching solution (10 mM ascorbate, 1 mM kojic acid and 1 mM EDTA in HBS). Cells were washed with quenching solution twice. Cells were washed twice with 10 mL ice-cold PBS, harvested by scraping, pelleted by centrifugation at 1,400 r.p.m. for 3 minutes, and either processed immediately or flash frozen in liquid nitrogen and stored at -80 °C before further analysis.

For the labeling of extracellular proteins by abmTYR with AxxP, MDA-MB-231 and HEK293T cells were washed twice with HBS and then treated with 75 nM of abmTYR and 50 μM of AxxP in HBS for 30 minutes. Cells were washed twice with ice-cold PBS, harvested by scraping, pelleted by centrifugation at 1,400 r.p.m. for 3 minutes, and either processed immediately or flash frozen in liquid nitrogen and stored at -80 °C before further analysis.

For the labeling of HER2-proximal proteins by ZHER-BmTYR and AxxP, MDA-MB-231 cells stably expressing GFP-HER2 were treated with 75 nM of ZHER-BmTYR in the culture medium at 37°C. After 30 minutes of incubation, the cells were washed with HBS three times. Next, the cells were incubated with 50 μM of AxxP in HBS for 30 minutes. For the spatial reference, MDA-MB-231 cells stably expressing GFP-HER2 were treated with 75 nM of untargeted BmTYR in the culture medium at 37°C. After 30 minutes of incubation, the cells were washed with HBS three times. Next, the cells were incubated with 50 μM of AxxP in HBS for 30 minutes. Cells were washed twice with ice-cold PBS, harvested by scraping, pelleted by centrifugation at 1,400 r.p.m. for 3 minutes, and either processed immediately or flash frozen in liquid nitrogen and stored at -80 °C before further analysis.

For the labeling of HER2-proximal proteins by ZHER-APEX2 and AxxP, MDA-MB-231 cells stably expressing GFP-HER2 were treated with 75 nM of ZHER-APEX2 in the culture medium at 37°C. For the spatial reference, MDA-MB-231 cells stably expressing GFP-HER2 were treated with 75 nM of untargeted APEX2 in the culture medium at 37°C. After 30 minutes of incubation, the cells were washed with HBS three times. Next, the cells were incubated with 50 μM of AxxP in HBS for 30 minutes. H_2_O_2_ (Sigma Aldrich) is added to a final concentration of 1 mM for 1 minute, with gentle agitation. The reaction was quenched by replacing the medium with an equal volume of quenching solution (10 mM ascorbate, 5 mM Trolox and 10 mM sodium azide in PBS). Cells were washed with quenching solution three times. Cells were washed twice with 10 mL ice-cold PBS, harvested by scraping, pelleted by centrifugation at 1,400 r.p.m. for 3 minutes, and either processed immediately or flash frozen in liquid nitrogen and stored at -80 °C before further analysis.

For all cell labeling experiments described above, the cells were lysed by resuspending in RIPA lysis buffer (50 mM Tris pH 8, 150 mM NaCl, 0.1% SDS, 0.5% sodium deoxycholate, 1% Triton X-100, 1× protease inhibitor cocktail (Sigma-Aldrich)) by gentle pipetting and incubating for 5 minutes at 4 °C. Notably, the RIPA buffer should be EDTA-free because EDTA interferes with the following copper-catalyzed click reaction. Lysates were clarified by centrifugation at 20,000 *g* for 10 minutes at 4 °C. Protein concentration in clarified lysates was estimated with Pierce BCA Protein Assay Kit (Thermo Fisher) and normalized to 2 mg/mL. 50 μL of lysates were reacted with 100 μM azide-Rhodamine, premixed 2-(4-((bis((1-tertbutyl-1H-1,2,3-triazol-4-yl)methyl)amino)methyl)-1H-1,2,3-triazol1-yl)-acetic acid (BTTAA)-CuSO_4_ complex (500 μM CuSO_4_, BTTAA:CuSO_4_ with a 2:1 molar ratio) and 2.5 mM freshly prepared sodium ascorbate for 2 hours at 25 °C. The excess reagents were removed by methanol and chloroform (4:1) precipitation and the fluorescent intensity was detected by in-gel fluorescence scanning.

### In-gel fluorescence scanning and Western blotting

For all in-gel fluorescence experiments, samples were resolved on a 10% SDS-PAGE gel and imaged by Tanon 5200multi. The gels were then stained by Coomassie brilliant blue to demonstrate equal loading. For all Western blots, samples were resolved on a 10% SDS-PAGE gel. After SDS-PAGE, the gels were transferred to a PVDF membrane, and then stained by Ponceau S (5 minutes in 0.1% (w/v) Ponceau S in 5% acetic acid/water). The blots were then blocked in 5% (w/v) BSA in TBS-T (Tris-buffered saline, 0.1% Tween 20) for at least 30 minutes at room temperature. For streptavidin blotting, the blots were stained with 0.3 μg/mL streptavidin-HRP in TBS-T for 1 hour at room temperature. For v5 blotting, the blots were stained with anti-v5 (1:3,000 dilution, catalog no. 460705, Thermo Fisher) for 2 hours at room temperature. After washing three times with TBS-T for 5 minutes each, the blots were stained with secondary antibodies in 5% BSA (w/v) in TBS-T for 1 hour in room temperature. The blots were washed three times with TBS-T for 5 minutes each time before to development with Clarity Western ECL Blotting Substrates (Thermo Fisher) and imaging on the Tanon 5200multi.

### Confocal fluorescence microscopy

To evaluate the impact of CuCl_2_ on BmTYR and organelle morphology, HEK 293T cells were plated on glass coverslips and transfected with BmTYR fusion construct of interest for 24 hours. Cells were washed twice with HBS, and then incubated with 5 μM CuCl_2_ in HBS for 1 hour, followed by the treatment of 50 μM AP for 30 minutes. The reaction was quenched by replacing the HBS with an equal volume of quenching solution (10 mM ascorbate, 1 mM kojic acid and 1 mM EDTA in HBS). Cells were washed with quenching solution twice. Cells were washed twice with 1 mL ice-cold PBS. Cells were fixed with cold methanol for 15 minutes at -20 °C and then washed three times with PBS. Alkyne labeled proteins were then reacted with 100 μM azide-Rhodamine, premixed 2-(4-((bis((1-tertbutyl-1H-1,2,3-triazol-4-yl)methyl)amino)methyl)-1H-1,2,3-triazol1-yl)-acetic acid (BTTAA)-CuSO_4_ complex (500 μM CuSO_4_, BTTAA:CuSO_4_ with a 2:1 molar ratio) and 2.5 mM freshly prepared sodium ascorbate in PBS for 1 hour at 25 °C. Cells were washed three times in PBS and blocked in 5% BSA dissolved in PBS for 1 hour. The coverslips were incubated with primary antibodies against v5 (1:200 dilution) and a marker of each organelle including anti-citrate synthetase (1:200 dilution, catalog no. BM5202, BOSTER), anti-Calnexin (1:200 dilution, catalog no.

A15631, ABclonal) and anti-GLUT1 (1:200 dilution, catalog no. ab15309, abcam) for 2 hours at room temperature or overnight at 4 °C. Coverslips were washed three times in PBS with 5 minutes for each wash then incubated in secondary antibodies conjugated to either AlexaFluor-488 (1:200 dilution, catalog no. 33206ES60, Yeasen) or 568 (1:200 dilution, catalog no. ab175470, abcam) and streptavidin conjugated to AlexaFluor647 (1:300 dilution, catalog no. 35104ES60, Yeasen) for 1 hour. Coverslips were washed five times in PBST (0.1% Tween-20 in PBS). Coverslips were incubated with DAPI (4’,6-diamidino-2-phenylindole, 1 μg/ml) for 10 minutes. Coverslips were washed three times, then mounted on glass slides.

To validate the cell surface labeling of abmTYR, HEK 293T and MDA-MB-231 cells were plated on glass coverslips. Cells were washed twice with HBS and treated with 75 nM of abmTYR and 500 μM of BP in HBS for 30 minutes. Cells were washed twice with 1 mL ice-cold PBS. Cells were fixed for 15 minutes with 4% paraformaldehyde at room temperature and washed in PBS three times. Cells were then permeabilized with 0.5% Triton X-100 (diluted in PBS) for 15 minutes at room temperature and washed in PBS three times. Cells were blocked in 5% BSA dissolved in PBS for 1 hour. The coverslips were incubated with primary antibodies against GLUT1 for 2 hours at room temperature or overnight at 4 °C. Coverslips were washed three times in PBS with 5 minutes for each wash then incubated in secondary antibodies conjugated to either AlexaFluor-488 or 568 and streptavidin conjugated to AlexaFluor647 for 1 hour. Coverslips were washed five times in PBST (0.1% Tween-20 in PBS). Coverslips were incubated with DAPI for 10 minutes. Coverslips were washed three times, then mounted on glass slides.

To validate the cell surface labeling of abmTYR with AxxP, HEK 293T and MDA-MB-231 cells were plated on glass coverslips. Cells were washed twice with HBS and treated with 75 nM of abmTYR and 50 μM of AxxP in HBS for 30 minutes. Cells were fixed with cold methanol for 15 minutes at -20 °C and then washed three times with PBS. Alkyne labeled proteins were then reacted with 100 μM azide-biotin, premixed 2-(4-((bis((1-tertbutyl-1H-1,2,3-triazol-4-yl)methyl)amino)methyl)-1H-1,2,3-triazol1-yl)-acetic acid (BTTAA)-CuSO_4_ complex (500 μM CuSO_4_, BTTAA:CuSO_4_ with a 2:1 molar ratio) and 2.5 mM freshly prepared sodium ascorbate in PBS for 1 hour at 25 °C. Cells were washed three times in PBS and blocked in 5% BSA dissolved in PBS for 1 hour. Coverslips were then incubated with primary antibodies against GLUT1 for 2 hours at room temperature or overnight at 4 °C. Coverslips were washed three times in PBS with 5 minutes for each wash then incubated in secondary antibodies conjugated to 568 and streptavidin conjugated to AlexaFluor647 for 1 hour. Coverslips were washed five times in PBST (0.1% Tween-20 in PBS). Coverslips were incubated with DAPI for 10 minutes. Coverslips were washed three times, then mounted on glass slides.

To validate HER2-selective cell surface labeling by ZHER-BmTYR and AxxP, HER2-positive and WT MDA cells were mixed and treated with 75 nM of ZHER-BmTYR in the culture medium at 37°C. After 30 minutes of incubation, the cells were washed with HBS three times. Next, the cells were incubated with 50 μM of AxxP in HBS for 30 minutes. After 30 minutes of incubation, the cells were washed twice with ice-cold PBS. The labeled proteins were then reacted with 100 μM azide-biotin, premixed 2-(4-((bis((1-tertbutyl-1H-1,2,3-triazol-4-yl)methyl)amino)methyl)-1H-1,2,3-triazol1-yl)-acetic acid (BTTAA)-CuSO_4_ complex (50 μM CuSO_4_, BTTAA:CuSO_4_ with a 6:1 molar ratio) and 2.5 mM freshly prepared sodium ascorbate in PBS for 10 minutes at 25 °C. Cells were washed three times in PBS and were fixed with 4% FPA for 15 minutes at 25 °C. Cells were washed three times in PBS and blocked in 5% BSA dissolved in PBS for 1 hour. Coverslips were then incubated with primary antibodies against GLUT1 for 2 hours at room temperature or overnight at 4 °C. Coverslips were washed three times in PBS (5 minutes for each wash), then incubated with secondary antibodies conjugated to 568 and streptavidin conjugated to AlexaFluor647 for 1 hour. Coverslips were washed five times in PBS. Coverslips were incubated with DAPI for 10 minutes. Coverslips were washed three times, then mounted on glass slides.

To validate HER2-selective cell surface labeling by ZHER-APEX2 and AxxP, HER2-positive and WT MDA cells were mixed and treated with 75 nM of ZHER-APEX2 in the culture medium at 37°C. After 30 minutes of incubation, the cells were washed with HBS three times. Next, the cells were incubated with 50 μM of AxxP in HBS for 30 minutes. H_2_O_2_ (Sigma Aldrich) is added to a final concentration of 1 mM for 1 minute, with gentle agitation. Then the cells were washed twice with ice-cold PBS. The labeled proteins were then reacted with 100 μM azide-biotin, premixed 2-(4-((bis((1-tertbutyl-1H-1,2,3-triazol-4-yl)methyl)amino)methyl)-1H-1,2,3-triazol1-yl)-acetic acid (BTTAA)-CuSO_4_ complex (50 μM CuSO_4_, BTTAA:CuSO_4_ with a 6:1 molar ratio) and 2.5 mM freshly prepared sodium ascorbate in PBS for 10 minutes at 25 °C. Cells were washed three times in PBS and were fixed with 4% FPA for 15 minutes at 25 °C. Cells were washed three times in PBS and blocked in 5% BSA dissolved in PBS for 1 hour. Coverslips were then incubated with primary antibodies against GLUT1 for 2 hours at room temperature or overnight at 4 °C. Coverslips were washed three times in PBS (5 minutes for each wash), then incubated with secondary antibodies conjugated to 568 and streptavidin conjugated to AlexaFluor647 for 1 hour. Coverslips were washed five times in PBS. Coverslips were incubated with DAPI for 10 minutes. Coverslips were washed three times, then mounted on glass slides.

To validate labeling in the intact mouse brain, adult mice were administered abmTYR and BP injections. One hour later, they were anesthetized with Avertin (500 mg/kg, intraperitoneal injection, Sigma-Aldrich). Following anesthesia, the mice were perfused with PBS followed by 4% paraformaldehyde (PFA) to fix the tissues. Brains were carefully dissected and post-fixed in 4% PFA overnight at 4°C. Subsequently, the brains were transferred to a 30% sucrose solution for overnight incubation at 4°C to facilitate cryoprotection and enhance tissue integrity. After two days, the dehydrated brain tissues were embedded in O.C.T. compound (Tissue-Tek) and rapidly frozen on dry ice. Serial coronal sections were then prepared at a thickness of 14 μm using a cryostat. For IF staining, the frozen brain sections were first oven-dried at 42°C for 30 minutes to enhance adherence to the slides. The sections were then rinsed and rehydrated with PBS. To enhance cell membrane permeability, the sections were treated with 0.3% Triton X-100 in PBS for 20 minutes at room temperature (RT). Blocking of nonspecific binding sites was achieved by incubating the sections with 2% normal goat serum in PBS for 1 hour at RT. Subsequently, the sections were incubated with Streptavidin-Alexa Fluor 647 dye (1:300 dilution, catalog no. 35104ES60, Yeasen) for 1 hour at RT. Afterward, the sections were counterstained with DAPI for 5 minutes to label cell nuclei. Finally, the slides were coverslipped using a suitable mounting medium and imaged under a Nikon A1 confocal microscope for detailed analysis of the stained brain sections.

### Cell viability assays

5.0×10^4^ HEK293T cells and 1.0×10^4^ MDA-MB-231cells were plated in 96-well plates with 100 μL of pre-warm complete DMEM medium per well (six technical replicates). For experiments that require overexpression, 0.1 μg NES-BmTYR plasmids were transfected for 24 h. Cells were labeled as described above. Cell viability was then detected by using MTS reagent (CellTiter 96® AQueous One Solution Cell Proliferation Assay, Promega). Cells were incubated in 100 μL serum-free medium (DMEM, Gibco) containing 1/6 of the MTS reagent at 37 °C for 30 minutes and the absorbance at 490 nm was measured with a microplate reader. For data presentation, the mean and standard deviation for the three biological replicates of each data point in a representative experiment were plotted in Prism 9 (Graphpad) and presented. The fitted curves were regressed by “Inhibitor vs Response” in Graphpad Prism software.

### Protein purification

ZHER-BmTYR, ZHER-abmTYR, BmTYR, APEX2 and ZHER-APEX2 were produced in Escherichia coli BL21 (DE3) cells with LB medium. Cells were grown at 37 ℃ until optical density at 600 nm (OD600) of 0.6-0.8. The temperature was then reduced to 16 ℃ and the cultures were grown for approximately 16 hours before harvesting. The cells were lysed in 20 mM Tris-HCl 8.0, 500 mM NaCl, 5% glycerol and 5 mM imidazole by sonication, the lysate was centrifuged at 20,000 g for 30 minutes at 4 ℃, the resulting supernatant was filtrated by 0.22 μm filter and then applied onto a 3-ml Ni NTA Beads 6FF column (Lablead). Bound protein was eluted with a gradient of imidazole from 50 to 500 mM. The fractions containing protein of interest was ultrafiltrated and changed solvent into PBS and stored at -80 ℃.

### Streptavidin enrichment

The products of click-labeled lysates or labeled whole-cell lysates were precipitated by methanol and chloroform (4:1). The nepheloid sediment were centrifuged for 5,000 g, 20 minutes at 4 ℃ and washed three times with 1 mL cold methanol. The proteomes were resuspended in 200 μL H_2_O with 1% SDS. The samples were sonicated and diluted to 0.2% SDS/RIPA.

For each sample, take 50 μL streptavidin magnetic beads (Thermo Fisher) and wash twice with 1 mL RIPA lysis buffer. Incubate the beads with 2 mg protein from each sample with an additional 1 mL RIPA lysis buffer at 4 °C overnight. Save the remaining material (store at −20 °C) from the whole-cell lysates for western blot analysis.

After enrichment, pellet the beads, using a magnetic rack, and collect the supernatant for western blot analysis. Wash the beads twice with 1 mL RIPA lysis buffer, once with 1 mL of 1 M KCl, once with 1mL of 0.1 M Na_2_CO_3_, once with 1 mL of 2 M urea in 10 mM Tris-HCl (pH 8.0), and twice with RIPA lysis buffer. Elute the enriched material from the beads by boiling each sample in 30 μL of 3× protein loading buffer supplemented with 2 mM biotin and 20 mM DTT at 95 °C for SDS-PAGE analysis or for LC-MS/MS analysis in the following steps.

### Dimethyl labeling-based quantitative proteomics

For all cell labeling experiments described above, the cells were lysed by resuspending in 200 μL of RIPA lysis buffer (50 mM Tris-HCl (Solarbio, Beijing, China), pH=8, 150 mM NaCl (Yuanye, Shanghai, China), 1% SDS (Sigma-Aldrich), 0.5% sodium deoxycholate (Macklin), 1% Triton X-100 (Biosharp)) containing an EDTA-free Pierce Halt™ protease inhibitor cocktail. After sonication on ice, the solution was supplemented with SDS-free RIPA buffer (50 mM Tris-HCl pH=8, 150 mM NaCl, 0.5% sodium deoxycholate, 1% Triton X-100) to a final volume of 1 mL. The cell lysates were centrifuged at 20,000 g for 30 minutes at 4 °C to remove the sediment. Notably, the RIPA buffer should be EDTA-free because EDTA interferes with the subsequent copper-catalyzed click reaction. The protein concentration in clarified lysates was estimated using the Pierce BCA Protein Assay Kit (Thermo Fisher) and normalized to 2 mg/mL. 50 μL of the 1 mL lysates were taken out for in-gel fluorescence analysis, and the remaining lysate was reacted with 100 μM azide-biotin, premixed with 2-(4-((bis((1-tert-butyl-1H-1,2,3-triazol-4-yl)methyl)amino)methyl)-1H-1,2,3-triazol-1-yl)-acetic acid (BTTAA) (Yuanye, Shanghai, China)-CuSO_4_ complex (500 μM CuSO_4_, BTTAA: CuSO_4_ with a 2:1 molar ratio), and 2.5 mM freshly prepared sodium ascorbate for 2 hours at 25 °C. The excess reagents were removed by methanol (TGREAG, Beijing, China) and chloroform (TGREAG, Beijing, China) (4:1) precipitation, followed by washing 3 times with 1 mL of cold methanol. The proteomes were resuspended in 0.5% SDS RIPA by sonication. Subsequently, SDS-free RIPA buffer was added to the solution to achieve a final SDS concentration of 0.1%. The proteome samples were then enriched using streptavidin beads as described above.

The enriched proteins with magnetic beads were washed twice by 1 mL PBS, the resulting beads were resuspended in 500 μL PBS with 6 M urea and 10 mM dithiothreitol (DTT) (Yuanye, Shanghai, China) at 35 ℃ for 30 minutes and alkylated by addition of 20 mM iodoacetamide (Yuanye, Shanghai, China) at 37 ℃ for 30 minutes in the dark. The beads were washed with 1 mL of 100 mM triethylammonium bicarbonate (TEAB) buffer (Sigma-Aldrich) in PBS. The beads were resuspended in 200 μL of 100 mM TEAB buffer with 2 M urea/PBS, 1 mM CaCl_2_ and 10 ng/μL trypsin (Meizhiyuan, Beijing, China). Trypsin digestion was performed at 37 ℃ on a thermomixer overnight.

Digested peptides released from the beads were collected in fresh microcentrifuge tubes. The beads were washed twice with 50 μL of 100 mM TEAB buffer, and the supernatant was combined with the peptide solution. For triple-dimethyl labeled samples, peptides were reacted with 12 μL of 4% D^13^CDO (Sigma-Aldrich), DCDO (Sigma-Aldrich), or HCHO (Sigma-Aldrich) respectively. For the ’light’ or ’medium’ labeled samples, 12 μL of 0.6 M NaBH_3_CN was added. For the ’heavy’ labeled samples, 12 μL of 0.6 M NaBD_3_CN was added to. For extracellular proteome profiling and HER2-neighboring proteome profiling, label swapping was performed across biological replicates. The labeling isotopes used in each experiment are shown in the table below. The reaction was incubated for 2 hours at room temperature, after which 48 μL of 1% ammonium hydroxide (Sigma-Aldrich) was added to terminate the reactions. The solutions were incubated for an additional 15 minutes at room temperature. Then, 24 μL of 2% formic acid (FA) (Sigma-Aldrich) was added to neutralize the remaining ammonium hydroxide. The peptides were fractionated by suspension and desalted using homemade C18 tips. The peptides were then sequentially fractionated using 6%, 12%, 15%, 18%, 21%, 25%, 30%, and 35% acetonitrile (ACN) (Thermo Fisher) with 10 mM ammonium hydrogen carbonate (ABC) (pH=10) (Solarbio, Beijing, China). The eluate containing the peptides was collected for LC-MS/MS analysis.

|  | Light | Medium | Heavy |
| --- | --- | --- | --- |
| extracellular<br>proteome_replicate_1 | +AxxP<br>+AbmTYR | +AxxP | +AbmTYR |
| extracellular<br>proteome_replicate_2 | +AbmTYR | +AxxP<br>+AbmTYR | +AxxP |
| extracellular<br>proteome_replicate_3 | +AxxP | +AbmTYR | +AxxP<br>+AbmTYR |
| ZHER-BmTYR<br>lableing_replicate_1 | +AxxP | +AxxP<br>+BmTYR | +AxxP<br>+ZHER-BmTYR |
| ZHER-BmTYR<br>lableing_replicate_2 | +AxxP<br>+BmTYR | +AxxP<br>+ZHER-BmTYR | +AxxP |
| ZHER-BmTYR<br>lableing_replicate_3 | +AxxP<br>+ZHER-BmTYR | +AxxP | +AxxP<br>+BmTYR |
| ZHER-APEX2<br>lableing_replicate_1 | +AxxP<br>+APEX2 | +AxxP<br>+APEX2<br>+H<sub>2</sub>O<sub>2</sub> | +AxxP<br>+ZHER-APEX2<br>+H<sub>2</sub>O<sub>2</sub> |
| ZHER-APEX2<br>lableing_replicate_2 | +AxxP<br>+APEX2<br>+H<sub>2</sub>O<sub>2</sub> | +AxxP<br>+ZHER-APEX2<br>+H<sub>2</sub>O<sub>2</sub> | +AxxP<br>+APEX2 |
| ZHER-APEX2<br>lableing_replicate_3 | +AxxP<br>+ZHER-APEX2<br>+H<sub>2</sub>O<sub>2</sub> | +AxxP<br>+APEX2 | +AxxP<br>+APEX2<br>+H<sub>2</sub>O<sub>2</sub> |

### Whole cell lysis proteomic profiling for TyroID labeling

Cells after TyroID labeling were lysed in 1 mL of TEAB buffer (Sigma) with 8 M urea (VETEC)/PBS containing the EDTA-free Pierce Halt™ protease inhibitor cocktail. After sonication on ice, the cell lysates were centrifuged at 20,000 g for 30 minutes at 4°C to remove the sediment. The protein concentration in the clarified lysates was estimated using the Pierce BCA Protein Assay Kit (Thermo Fisher) and normalized to 6 mg/mL. 100 μL lysates were subjected to the following experiments. 1 μL of 1 M DTT (Yuanye, Shanghai, China) was added to each sample and incubated at 35°C for 30 minutes. The proteins were then alkylated by adding 3 μL of 1 M iodoacetamide (Sigma) and incubating at 37°C for 30 minutes in the dark. The reaction was quenched by adding 0.5 μL of 1 M DTT and incubating at 35°C for 30 minutes. The solutions were diluted in 400 μL of TEAB buffer with 2 M urea/PBS, 1 mM CaCl_2_, and 40 ng/μL trypsin (Meizhiyuan, Beijing, China). Trypsin digestion was performed on a thermomixer at 37°C overnight.

30 μL of digested samples were diluted with TEAB buffer to a final volume of 500 μL. For triple-dimethyl labeled samples, peptides were reacted with 20 μL of 4% D^13^CDO (Sigma-Aldrich), DCDO (Sigma-Aldrich), or HCHO (Sigma-Aldrich), respectively. For the ’light’ or ’medium’ labeled samples, 20 μL of 0.6 M NaBH_3_CN was added. For the ’heavy’ labeled samples, 20 μL of 0.6 M NaBD_3_CN was added. The labeling isotopes used in each experiment are shown in the table below. The reaction mixtures were incubated for 2 hours at room temperature, after which 48 μL of 1% ammonium hydroxide (Sigma-Aldrich) was added to terminate the reactions. The solutions were incubated for an additional 15 minutes at room temperature. Then, 24 μL of 2% formic acid (FA) (Sigma-Aldrich) was added to neutralize the remaining ammonium hydroxide. The peptides were fractionated by suspension and desalted using homemade C18 tips. The peptides were then sequentially fractionated using 6%, 12%, 15%, 18%, 21%, 25%, 30%, and 35% acetonitrile (ACN) (Thermo Fisher) with 10 mM ammonium hydrogen carbonate (ABC) (pH=10) (Solarbio, Beijing, China). The eluates containing the peptides were collected for LC-MS/MS analysis.

|  | Light | Medium | Heavy |
| --- | --- | --- | --- |
| Whole_lysis_replicate_1 | +AxxP | +AbmTYR | +AxxP<br>+AbmTYR |
| Whole_lysis_replicate_2 | +AbmTYR | +AxxP<br>+AbmTYR | +AxxP |
| Whole_lysis_replicate_3 | +AxxP<br>+AbmTYR | +AxxP | +AbmTYR |

### Preparation of acid-cleavable azide-functionalized Beads

The preparation of agarose beads functionalized with azide groups and acid-cleavable linkers was carried out by following a previous study^49^. The NHS beads (Cytiva) were sequentially washed twice with 10 mL of 1 mM cold HCl and 10 mL of 0.1% Triton (Biosharp) /PBS. The beads were then resuspended in 10 mL of 0.1% Triton/PBS. For each experiment, 2 mL of solid beads and 200 μL of 100 mM CY75 stock solution (gift from chu wang lab) were used. After incubation for 6 hours at room temperature or overnight at 4 °C, the beads were further incubated with blocking buffer (100 mM Tris-HCl (Solarbio, Beijing, China), pH 8.0, 150 mM NaCl, 100 mM ethanolamine) to block any remaining NHS groups. The blocking time ranged from 6 to 12 hours, depending on the reaction temperature. The blocked beads were washed with 10 mL of PBS for five times and once with 10 mL of 30% glycerol (Solarbio, Beijing, China) /PBS. Finally, the beads were resuspended in 30% glycerol/PBS, aliquoted into 40 equal portions, and stored at −20 °C until use. Frequent freezing and thawing of the CY75-functionalized beads should be avoided.

### SuperTOP-ABPP

To map the modification sites of BmTYR labeling in the mitochondrial matrix, HEK293T cells were transiently transfected with mito-BmTYR for 24 hours. Cells were washed twice with HBS, and then incubated with 5 μM CuCl_2_ in HBS for 1 hour, followed by the treatment of 50 μM alkyne-phenol for 30 minutes. The reaction was quenched by replacing the HBS with an equal volume of quenching solution (10 mM ascorbate, 1 mM kojic acid and 1 mM EDTA in HBS). Cells were washed with quenching solution twice. Cells were washed twice with 10 mL ice-cold PBS, harvested by scraping, pelleted by centrifugation at 1,400 r.p.m. for 3 minutes, and either processed immediately or flash frozen in liquid nitrogen and stored at -80 °C before further analysis.

To map the modification sites of APEX2 labeling in the mitochondrial matrix, HEK293T cells were transiently transfected with mito-APEX2 for 24 hours. APEX2 labeling was initiated by changing to fresh medium containing 50 μM alkyne-phenol and incubating at 37 °C under 5% CO_2_ for 30 minutes. H_2_O_2_ (Sigma Aldrich) is added to a final concentration of 1 mM for 1 minute, with gentle agitation. The reaction was quenched by replacing the medium with an equal volume of quenching solution (10 mM ascorbate, 5 mM Trolox and 10 mM sodium azide in PBS). Cells were washed with quenching solution three times to remove excess probe. Cells were washed twice with 10 mL ice-cold PBS, harvested by scraping, pelleted by centrifugation at 1,400 r.p.m. for 3 minutes, and either processed immediately or flash frozen in liquid nitrogen and stored at -80°C before further analysis.

To map the modification sites of AxxP by abmTYR, MDA-MB-231 and HEK293T cells were washed twice with HBS and then treated with 75 nM of abmTYR and 50 μM of AxxP in HBS for 30 minutes. Cells were washed twice with ice-cold PBS, harvested by scraping, pelleted by centrifugation at 1,400 r.p.m. for 3 minutes, and either processed immediately or flash frozen in liquid nitrogen and stored at - 80 °C before further analysis.

For cell labeling experiments described above, the cells were lysed by resuspending in 700 μL of PBS with 0.1% Triton X-100 (Biosharp) containing an EDTA-free Pierce Halt™ protease inhibitor cocktail. After sonication on ice, the cell lysates were centrifuged at 20,000 g for 30 minutes at 4°C to remove the sediment. The protein concentration in clarified lysates was estimated using the Pierce BCA Protein Assay Kit (Thermo Fisher) on a microplate and adjusted to 2 mg/mL. The lysates were precipitated using the methanol (TGREAG, Beijing, China) and chloroform mixture (4:1) (TGREAG, Beijing, China). The protein pellets were then resuspended in 0.4% SDS/PBS. The proteins were reduced with 10 mM dithiothreitol (DTT) (Yuanye, Shanghai, China) at 35°C for 30 minutes and alkylated with 20 mM iodoacetamide (IAA) (Sigma-Aldrich) at 37°C for 30 minutes in the dark on a thermomixer. The alkylated proteins were precipitated again with methanol-chloroform (4:1), and the protein pellets were washed three times with ice-cold methanol. The proteins were then resuspended in PBS buffer with ultrasonication. The protein was digested with 10 ng/μL of trypsin (Meizhiyuan, Beijing, China) for 16 hours at 37°C on a thermomixer. After digestion, the peptides were mixed with acid-cleavable azide-functionalized beads. A final concentration of 1 mM CuSO_4_ and 2 mM 2-(4-((bis((1-tert-butyl-1H-1,2,3-triazol-4-yl)methyl)amino)methyl)-1H-1,2,3-triazol-1-yl)-acetic acid (BTTAA) (Yuanye, Shanghai, China) was added, along with a final concentration of 0.5% sodium ascorbate (J&K Scientific, Beijing, China) to reduce Cu^2+^. The click reaction was carried out at 29°C for 2 hours on a thermomixer. The beads were washed sequentially with 1 mL of 8 M urea (VETEC), twice with 1 mL PBS, and five times with H_2_O. The washing method involved centrifuging the peptide sample with beads at 2000 g for 5 minutes, removing the supernatant, and repeating the process. Peptides were subjected to on-beads cleavage using 200 μL of 2% FA (Thermo Fisher) twice for 30 minutes on a thermomixer. The peptide eluates were collected by centrifuging at 2000 g for 5 minutes for LC-MS/MS analysis.

### LC-MS/MS

Samples were analyzed by LC-MS/MS on Q Exactive-plus series Orbitrap mass spectrometers (Thermo Fisher Scientific) coupled with Vanquish™ Neo UHPLC. Mobile phase A was 0.1% FA in H_2_O, and mobile phase B was 0.1% FA in ACN. The flow rate was 3 μL/ minutes for loading and 0.3 μL/ minutes for eluting. Labeled peptide samples were loaded onto a 100 μm fused silica column packed with 15 cm × 3 μm C18 resin. Under the positive-ion mode, full-scan mass spectra were acquired over the m/z range from 350 to 1800 using the Orbitrap mass analyzer with mass resolution of 70 000. MS/MS fragmentation is performed in a data-dependent mode, of which the TOP 20 most intense ions are selected for MS2 analysis a resolution of 17 500 using collision mode of HCD. Other important parameters: isolation window, 2.0 m/z units; default charge, 2+; normalized collision energy, 28%; maximum IT, 50 ms; and dynamic exclusion, 20.0 s.

### Data analysis

For dimethyl labeling data analysis, the MS/MS data were processed with the MaxQuant software (v2.4.11.0), carbamidomethyl cysteine was set as fixed modification, and methionine oxidation and acetyl N-terminal were set as variable modifications. DimethLys0 and DimethNter0 were enabled in the light labels, DimethLys4 and DimethNter4 were enabled in the medium labels. DimethLys8 and DimethNter8 were enabled in the heavy labels. The minimal peptide length was set to 7 amino acids, and the maximal mass tolerance was set to 20 ppm and 4.5 ppm in the first search and main search. The false discovery rate (FDR) was set to 1%. Proteins with at least two unique peptides were reported as identified proteins. For protein quantification, the functions of “re-quantify” and “match between runs” were enabled, and proteins with at least two quantified peptides (at least one unique peptide) were defined as quantified proteins.

After the program finished, dimethyl labeling data analysis depended on the output file “combined-txt-proteinGroups.txt”, including Ratio H/L, Ratio H/M, and so on. Based on the above information, we can sort out the dimethyl labeling quantification and obtain the corresponding protein information.

For the site data analysis, the MS/MS data were processed with the Fragpipe software (v21.1) integrated with MSFragger 4.0, IonQuant 1.10.12, Philosopher 5.1.0. The mass offset search strategy was used, and loaded the ’Mass-Offset-CommonPTMs’ workflow, allowing selected mass differences within 20ppm tolerance on target peptide match. To perform the restricted offset search, enter the mass offset masses of interest in the mass offsets box on the MSFragger tab, such as MDA-MB-231 cells treated with 75 nM of abmTYR and 50 μM of AxxP for 30 minutes, the mass offset masses were set 534.26896 and 536.28461. Next, enter the amino acid site(s) at which to allow mass offsets in the “Restrict delta mass to” box on the MSFragger tab, such as “CHK”, on behalf of cysteine, histidine, and lysine. At the same time, because MDA-MB-231 cells are of human origin, the human database should be selected for search and follow the prompts to use the default settings. Reviewed human sequences with common contaminants were downloaded from Uniprot on Oct.26, 2023; with 42460 proteins. In the MSFragger tab, methionine oxidation was set as a variable modification in the Variable Modifications section, and cysteine carbamidomethylation should be set as a variable mod instead of a fixed mod. In general, parameter settings are also common, fragment mass tolerance was set to 20 ppm, and number of top-N peaks used for matching was set to 100, and the precursor tolerance range was set to -10 to 10 ppm. The peptide mass range was set from 500 to 5,000 Da, and peptides with length 7 to 50 residues were allowed. One missed cleavage per peptide was allowed.

After the program finished, site data analysis depended on “output files-psm.tsv”, including Observed Modifications, MSFragger Localization, Protein ID, and so on. Based on the above information, we can sort out the target modification sites and their amino acids and peptides, and obtain the corresponding protein information. After that, we can perform a series of mapping of subsequently modified peptides and search for secondary spectra.

### Flow cytometry

Cells were incubated with ZHER-BmTYR in the culture medium at 37°C. After 30 minutes of incubation, the cells were washed with HBS three times. Next, the cells were incubated with 500 μM BP in HBS for 20 minutes. Cells were washed twice with PBS and dissociated with trypsin. The resulting cell suspension was stained with 3 μg/ml streptavidin-AF647 for 30 minutes in PBS. After three more washes with PBS, the cells were subjected to flow cytometry analysis.

Flow cytometry was performed with a BD FACSymphony A5SE Flow Cytometer. We used an SSC-A over FSC-A graph to discriminate cells from debris. Flow cytometry results were analyzed using BD FlowJo v10.8.1.

### Co-immunoprecipitation

To validate HER2-proximal proteins, 1 × 10⁶ MDA-MB-231 cells were seeded in 10 cm dishes and cultured overnight. The cells were transiently transfected with N-terminal FLAG-tagged HER2. After 8 hours of transfection, the medium was replaced with fresh medium. The cells were collected 24 hours post-transfection, washed three times with PBS by centrifugation at 3000 rpm for 4 minutes. The cells were then lysed in 500 μL of BC100 buffer (100 mM NaCl (Yuanye, Shanghai, China), 20 mM Tris (Solarbio, Beijing, China), pH=7.3, 20% glycerol (Solarbio, Beijing, China), 0.2% NP-40 (Solarbio, Beijing, China)) containing the EDTA-free Pierce Halt™ protease inhibitor cocktail. A fraction of the lysates (5% or 10%) was collected as input samples. The remaining lysates were incubated with 20 μL of anti-FLAG M2 affinity gel beads (1:1) (Sigma-Aldrich) at 4 °C overnight. The beads were then washed five times with BC100 buffer and mixed with 5×SDS-PAGE loading buffer, followed by heating at 95 °C. The supernatant was separated by SDS-PAGE and immunoblotted with indicated antibodies.

For the immunoprecipitation of endogenous HER2, SK-BR3 cells were collected, washed three times with PBS by centrifugation at 3000 rpm for 4 minutes. The cells were lysed in 1 mL of BC100 buffer. 10% of the cell lysates were collected as input samples, and the remaining lysates were divided into two aliquots. One aliquot was incubated with 1 μg of anti-HER2 antibody, while the other was incubated with normal rabbit IgG (Santa Cruz Biotechnology) at 4 °C overnight. Protein A/G agarose beads (Santa Cruz Biotechnology) were added to the lysates and incubated for 2 hours at 4 °C. The agarose beads were washed five times with BC100 buffer and mixed with 5×SDS-PAGE loading buffer, followed by heating at 95 °C. The supernatant was separated by SDS-PAGE and immunoblotted with indicated antibodies.

### Animal experiments

For xenograft and intravenous mouse model, male BALB/c nude mice aged 3-4 weeks and female C57BL/6J mice aged 6-7 weeks were obtained from Beijing Vital River Laboratory Animal Technology Co., Ltd (Beijing, China). The animals were housed in pressurized, individually ventilated cages (PIV/IVC) and maintained under specific-pathogen-free conditions, with free access to food and water in a controlled 12-hour light/dark cycle. All animal experiments were conducted in accordance with the ethical standards and approved by the Institutional Animal Care and Use Committees of Tsinghua University (Beijing, China).

For intracranial surgery experiment, female C57BL/6J mice aged 7-8 weeks were obtained from the Laboratory Animal Center of the Institute of Genetics and Developmental Biology, Chinese Academy of Sciences (Beijing, China). The animals were housed in pressurized, individually ventilated cages (PIV/IVC) and maintained under specific-pathogen-free conditions, with free access to food and water in a controlled 12-hour light/dark cycle. All animal experiments were conducted in accordance with the ethical standards and approved by the Animal Care and Use Committee of the Institute of Genetics and Developmental Biology, Chinese Academy of Sciences (Beijing, China).

### Xenograft Mouse Model

MDA-231 cells were resuspended in PBS with the density of 3 × 10^7^ cells per ml, and were injected into the flank of hind legs of BALB/c nude mice in a volume of 0.1 ml on both sides through s.c.. 1 week later, the tumor volume was about 100 mm^3^ measured by caliper, then mice were anesthetized and ZHER-BmTYR (50 μM solution in PBS) was multi-point injected intratumorally in a total volume of 0.1 ml into the right tumor while PBS was injected intratumorally into the left tumor in the same way. 15 minutes after enzyme injection, AxxP (10 mM solution in PBS) was injected intratumorally in a total volume of 0.05 ml into both sides tumor. After 12 hours, repeat previous injection and mice were euthanized and tumors were removed 30 minutes after the second injection. The maximum tumor size approved by IACUC protocol was 2 cm in diameter and this was not exceeded in all experiments.

### Intravenously Mouse Model

C57BL/6J mice were anesthetized with isoflurane inhalation, 2μL AxxP (500 mM solution in DMSO) was added into 0.1 ml abmTYR (120 μM solution in PBS) and mixed mildly. The mixed solution was immediately injected through i.v. in a volume of 0.1 ml. Blood plasma were obtained 1 or 12 hours after injection.

### Mice Intracranial Surgery

Mice were anesthetized with isoflurane inhalation and placed in a stereotaxic frame (RWD Life Science, Shenzhen, China). Intracranial injections were performed using a glass microelectrode needle (RWD Life Science, Shenzhen, China). For the injection, a mixture of BP (25 nl of a 500 mM BP solution in dimethylsulfoxide (DMSO, Sigma-Aldrich) and 525nl Phosphate Buffered Saline (PBS)) was administered into the right HP, while a combination of BP (25nl of a 500 mM solution in DMSO), 25nl PBS, and abmTYR (500nl of a 120 μM solution in PBS) was injected into the left HP. The stereotactic coordinates used were 1.8 mm posterior to the bregma (A.P: -1.8 mm), 1.2 mm lateral to the midline (M.L: ±1.2 mm), and 1.5 mm deep (D.V: -1.5 mm). Following the injection, the needle was left in place for 10 minutes to minimize the backflow of the injected solutions.

### Orthotopic transplanted mouse model

GL261 glioma cells were dissociated by 0.25% trypsin-EDTA. Single-cell suspensions were prepared in sterile Hanks Balanced Salt Solution (HBSS, Gibco) immediately before the stereotaxically injection procedure. 2x10^4^ cells in 2µL HBSS were injected into the striatum of female C57BL/6J mice at 7-8 weeks of age using a 10μL micro-syringe (Hamilton). Stereotactic coordinates used were 0.5 mm anterior to the bregma, 2.5 mm lateral to the midline, and 2.5 mm deep.

Statistics and reproducibility. For STRING analysis, we selected a medium evidence level for the edges to represent the network. For GO analysis, we used DAVID bioinformatics for online analysis and the human proteome as background. Three replicates were performed for all experiments with similar results. For comparison between two groups, P values were determined using two-tailed Student’s t-tests, *P < 0.05; **P < 0.01; ***P < 0.001; N.S. not significant. Error bars represent means ±SD

## Notes

### Competing Interest Statement

The authors have declared no competing interest.

