## Supporting Information for "Spatiotemporally-resolved mapping of extracellular proteomes via *in vivo*-compatible TyroID"

[1]School of Pharmaceutical Sciences, Tsinghua University, Beijing, 100084, China. [2]Tsinghua-Peking Center for Life Sciences, Tsinghua University, Beijing, 100084, China. [3]MOE Key Laboratory of Bioorganic Phosphorus Chemistry & Chemical Biology, Tsinghua University, Beijing, 100084, China. [4]The State Key Laboratory of Membrane Biology, Tsinghua University, Beijing, 100084, China. [5]Beijing Frontier Research Center for Biological Structure, Tsinghua University, Beijing, 100084, China. [6]State Key Laboratory of Molecular Developmental Biology, Institute of Genetics and Developmental Biology, Chinese Academy of Sciences, Beijing, 100101, China.

### Supplementary Figures

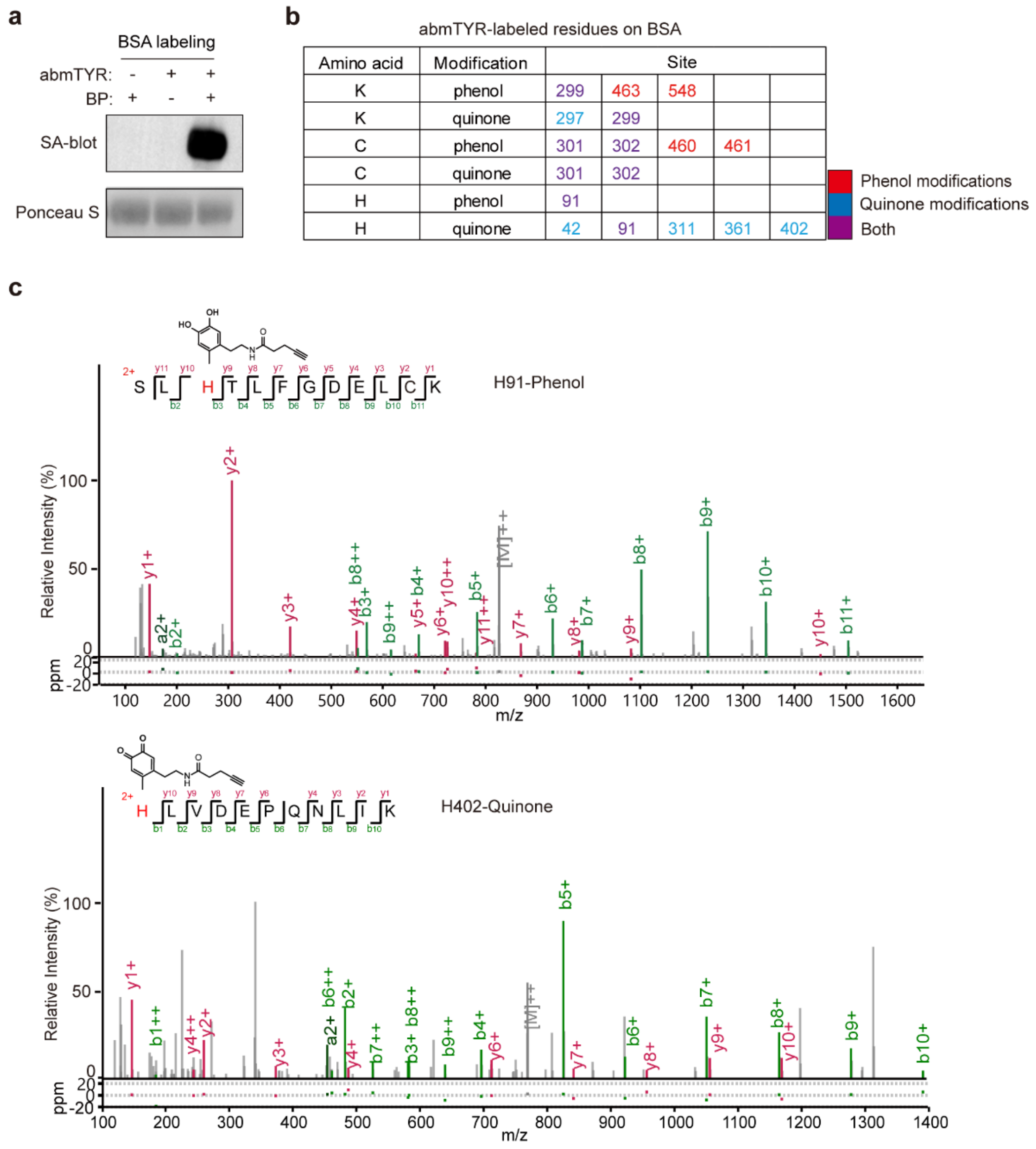


| Amino acid | Modification | Site | | | | |
|---|---|---|---|---|---|---|
| K | phenol | 299 | 463 | 548 | | |
| K | quinone | 297 | 299 | | | |
| C | phenol | 301 | 302 | 460 | 461 | |
| C | quinone | 301 | 302 | | | |
| H | phenol | 91 | | | | |
| H | quinone | 42 | 91 | 311 | 361 | 402 |

**Figure S1. Validation of tyrosinase-based labeling on BSA. a.** The labeling of bovine serum albumin (BSA) by abmTYR and AP *in vitro*. Recombinant BSA was treated with 75 nM abmTYR and 50 µM BP for 30 minutes, followed by streptavidin blotting. **b.** Summary of abmTYR-labeled residues on BSA. **c.** Tandem MS spectra of abmTYR-labeled peptides from BSA, including phenol and quinone modifications.

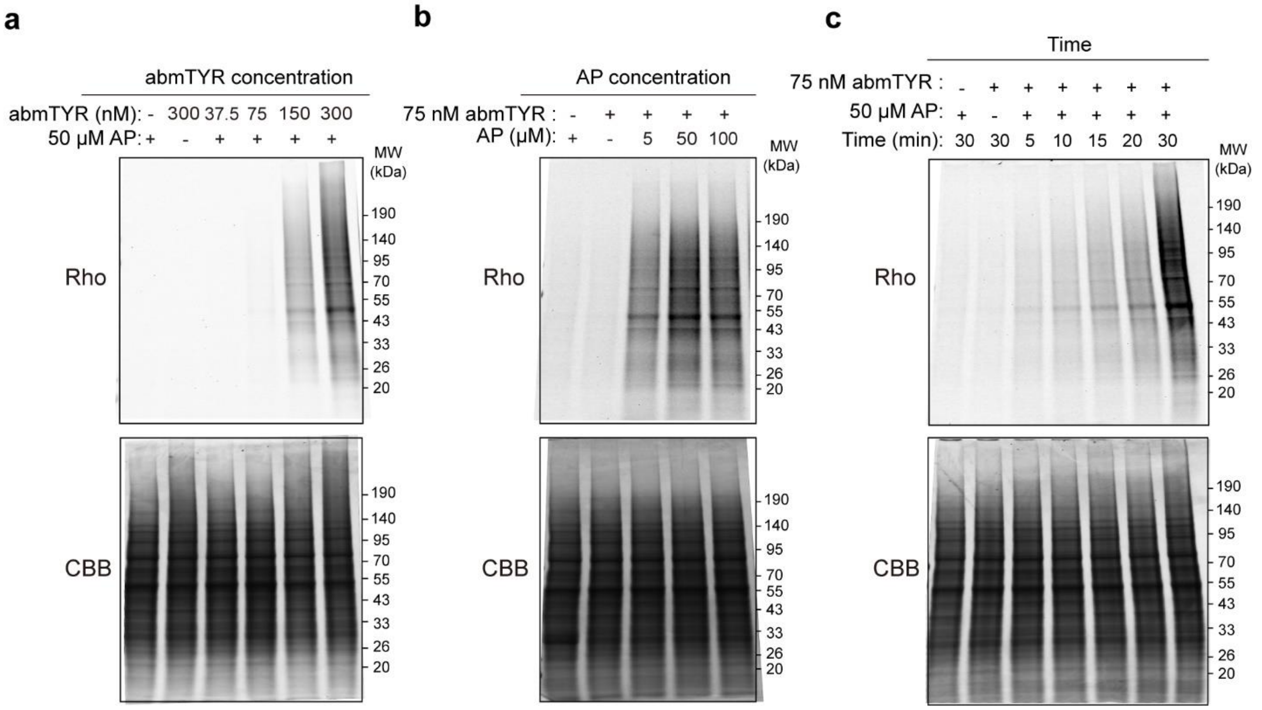


**Figure S2. Validation of tyrosinase-based protein labeling in cell lysates. a.** The impact of enzyme concentration on abmTYR labeling in cell lysates. HEK293T cell lysates were treated with various concentrations of abmTYR and 50 μM AP for 30 minutes, followed by click reaction with azide-Rhodamine and in-gel fluorescence scanning. **b.** The impact of probe concentration on abmTYR labeling in cell lysates. HEK293T cell lysates were treated with 75 nM abmTYR and various concentrations of AP for 30 minutes, followed by click reaction with azide-Rhodamine and in-gel fluorescence scanning. **c.** Time-dependent abmTYR labeling in cell lysates. HEK293T cell lysates were treated with 75 nM abmTYR and 50 μM AP for various duration of time. For **a-c**, cell lysates were clicked with azide-Rhodamine and analyzed by in-gel fluorescence scanning.

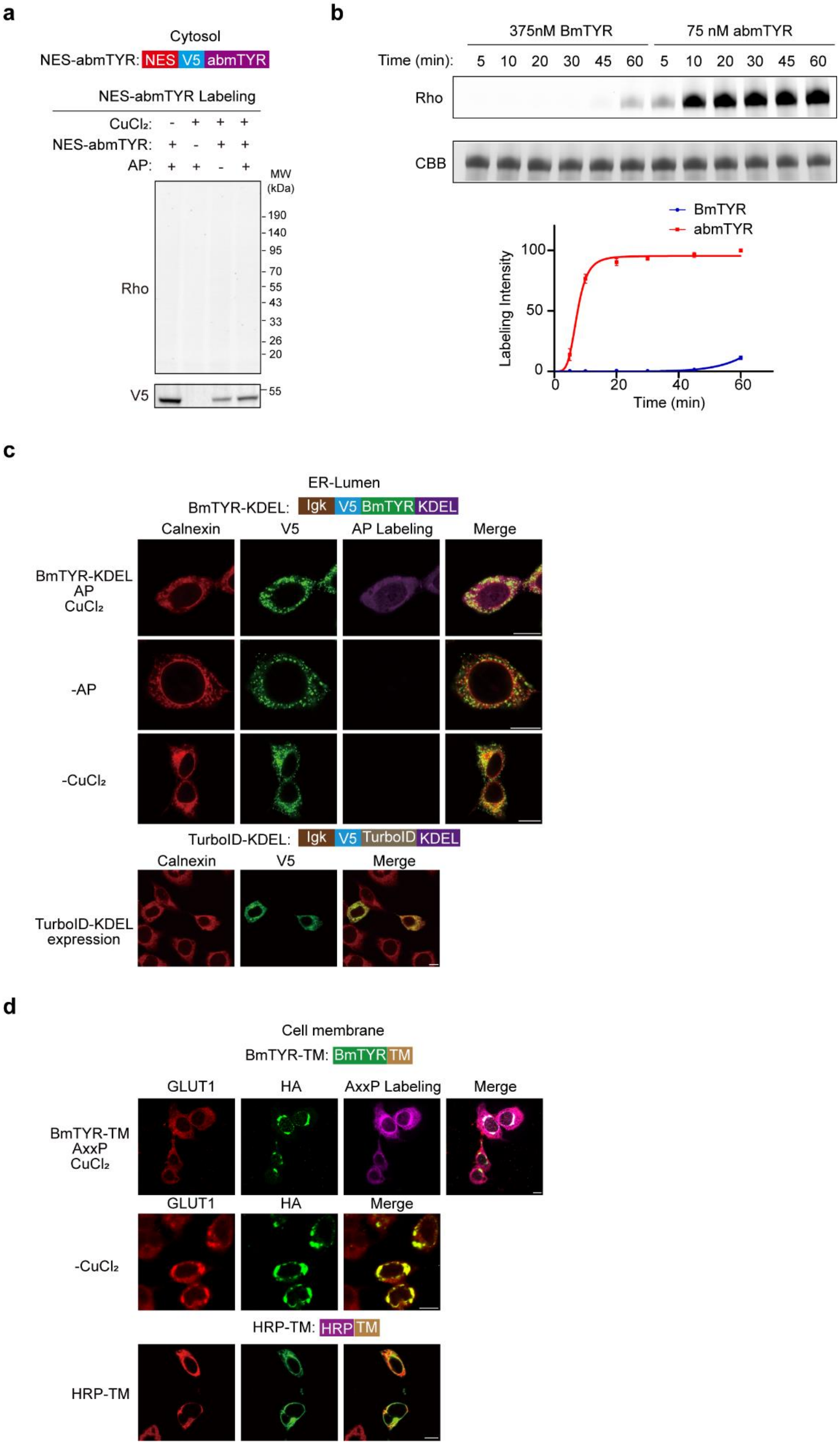
a
Cytosol
NES-abmTYR: NES V5 abmTYR
NES-abmTYR Labeling
CuCl₂: - + + +
NES-abmTYR: + - + +
AP: + + - +
MW (kDa)
190
140
95
70
55
43
33
26
20
Rho
V5
55
b
375nM BmTYR
75 nM abmTYR
Time (min): 5 10 20 30 45 60 5 10 20 30 45 60
Rho
CBB
BmTYR
abmTYR
Labeling Intensity
100
50
0
0 20 40 60
Time (min)
c
ER-Lumen
BmTYR-KDEL: Igk V5 BmTYR KDEL
Calnexin
V5
AP Labeling
Merge
BmTYR-KDEL
AP
CuCl₂
-AP
-CuCl₂
TurboID-KDEL: Igk V5 TurboID KDEL
Calnexin
V5
Merge
TurboID-KDEL expression
d
Cell membrane
BmTYR-TM: BmTYR TM
GLUT1
HA
AxxP Labeling
Merge
BmTYR-TM
AxxP
CuCl₂
GLUT1
HA
Merge
-CuCl₂
HRP-TM: HRP TM
HRP-TM

**Figure S3. Evaluation of the labeling by genetically-encode tyrosinases in living cells. a.** The expression and labeling of the H subunit from abmTYR in HEK293T cells. HEK293T cells were transfected with the H subunit from abmTYR for 24 hours, followed by the treatment of 5 μM $CuCl_2$ and 50 μM AP. The cells were lysed and subjected to click-based in-gel fluorescence detection. V5 indicates the expression of BmTYR. **b.** Comparison of labeling kinetics between bacterial tyrosinase (BmTYR) and mushroom tyrosinase (abmTYR). BSA was incubated with 375 nM of BmTYR or 75 nM of abmTYR and 50 μM of alkyne-phenol for different periods. The labeled BSA was reacted with azide-Rhodamine via click chemistry, followed by in-gel fluorescence detection. Quantification of the labeling intensity from three replicates was shown below. The error bars show mean ± SD. **c.** Confocal fluorescence imaging of BmTYR-KDEL and its impact on ER morphology. HEK293T cells were transfected with BmTYR-KDEL for 24 hours, treated with 5 μM $CuCl_2$ for 1 hour, followed by the treatment of 50 μM AP for 30 minutes. The cells were fixed and click with azide-biotin after cell fixation to visualize the labeled proteins. Streptavidin-AF647 indicates biotin labeling and Calnexin indicates the ER. Scale bars, 10 μm. **d.** Confocal fluorescence imaging of BmTYR-TM and its impact on plasma membrane. HEK293T cells were transfected with BmTYR-TM for 24 hours, treated with 5 μM $CuCl_2$ for 1 hour, followed by the treatment of 50 μM AP for 30 minutes. The cells were fixed and click with azide-biotin after cell fixation to visualize the labeled proteins. Streptavidin-AF647 indicates biotin labeling and GLUT1 indicates the plasma membrane. Scale bars, 10 μm.

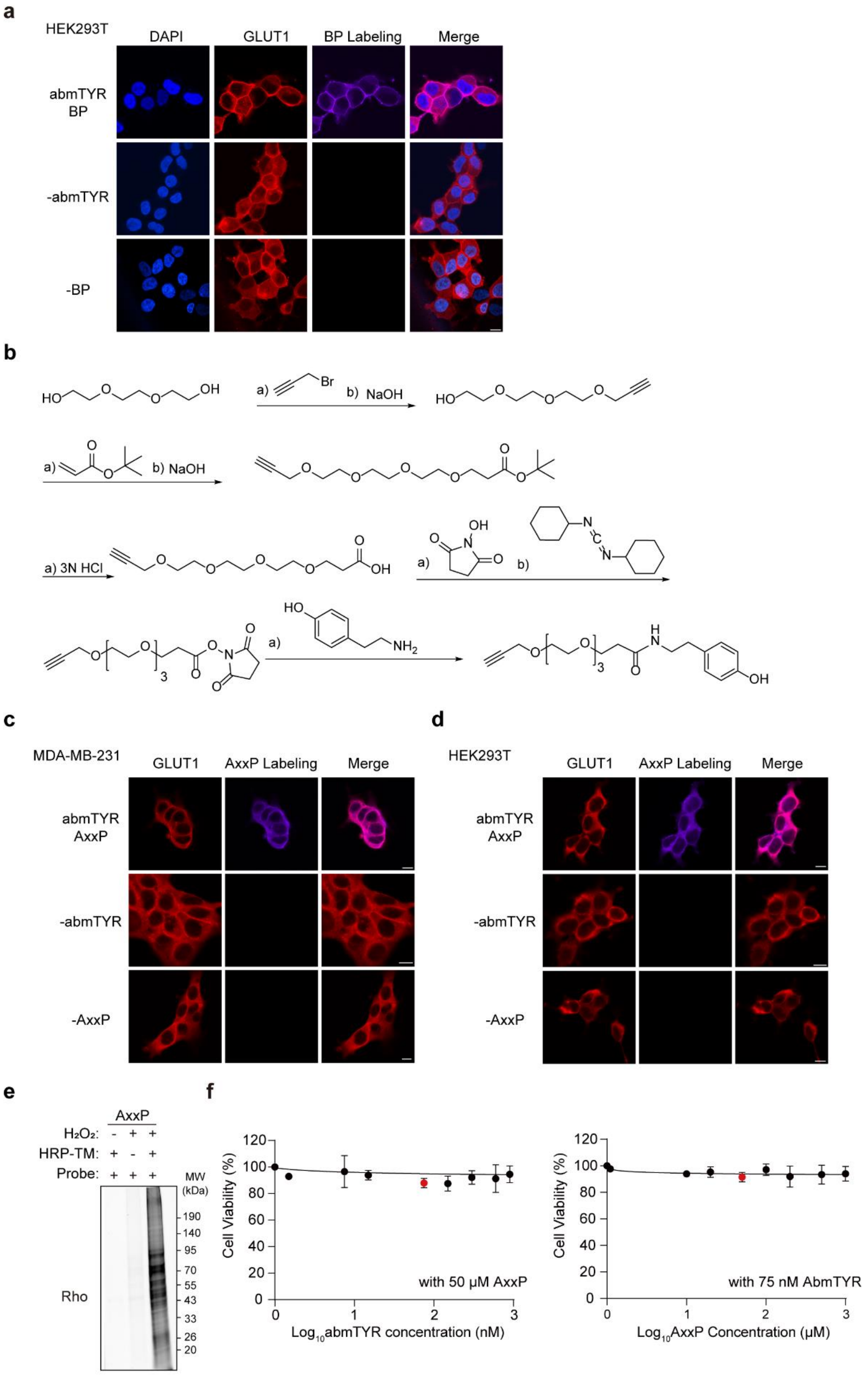
a
HEK293T
DAPI
GLUT1
BP Labeling
Merge
abmTYR
BP
-abmTYR
-BP
b
HO
O
O
OH
a)
Br
b) NaOH
HO
O
O
O
a)
O
O
b) NaOH
O
O
O
O
O
O
a) 3N HCl
O
O
O
O
O
OH
OH
N
O
O
a)
b)
N
C
N
O
O
3
O
O
N
O
O
HO
a)
NH2
O
O
3
H
N
O
OH
c
MDA-MB-231
GLUT1
AxxP Labeling
Merge
abmTYR
AxxP
-abmTYR
-AxxP
d
HEK293T
GLUT1
AxxP Labeling
Merge
abmTYR
AxxP
-abmTYR
-AxxP
e
AxxP
H₂O₂: - + +
HRP-TM: + - +
Probe: + + +
MW
(kDa)
190
140
95
70
55
43
33
26
20
Rho
f
Cell Viability (%)
120
100
80
60
40
20
0
with 50 μM AxxP
0
1
2
3
Log10abmTYR concentration (nM)
Cell Viability (%)
with 75 nM AbmTYR
Log10AxxP Concentration (μM)

**Figure S4. Development of membrane-impermeable alkyne-phenol for extracellular labeling by recombinant tyrosinases.** **a.** Confocal fluorescence imaging of cell surface proteins labeled by recombinant abmTYR with BP in HEK293T cells. Streptavidin-AF647 indicates biotin labeling, GLUT1 and DAPI represents the plasma membrane and nucleus respectively. Scale bars, 10 μm. **b.** The synthesis route of membrane-impermeable alkyne-phenol (AxxP). **c, d.** Confocal fluorescence imaging of cell surface proteins labeled by recombinant abmTYR with AxxP in MDA-MB-231 (**b**) and HEK293T (**c**) cells, click with azide-biotin after cell fixation to visualize the labeled proteins. Streptavidin-AF647 indicates biotin labeling, GLUT1 represents the plasma membrane. Scale bars, 10 μm. **e.** Evaluation of HRP labeling with AxxP. HEK293T cells were transfected with HRP-TM and incubated with 500 μM AxxP for 30 minutes, followed by 1 minute of $H_2O_2$ treatment. The cells were lysed and subjected to click-based in-gel fluorescence detection. **f.** Cell viability quantification at different doses of abmTYR (left) and AxxP (right). Three biological replicates were performed, and the error bars show mean ± SD.

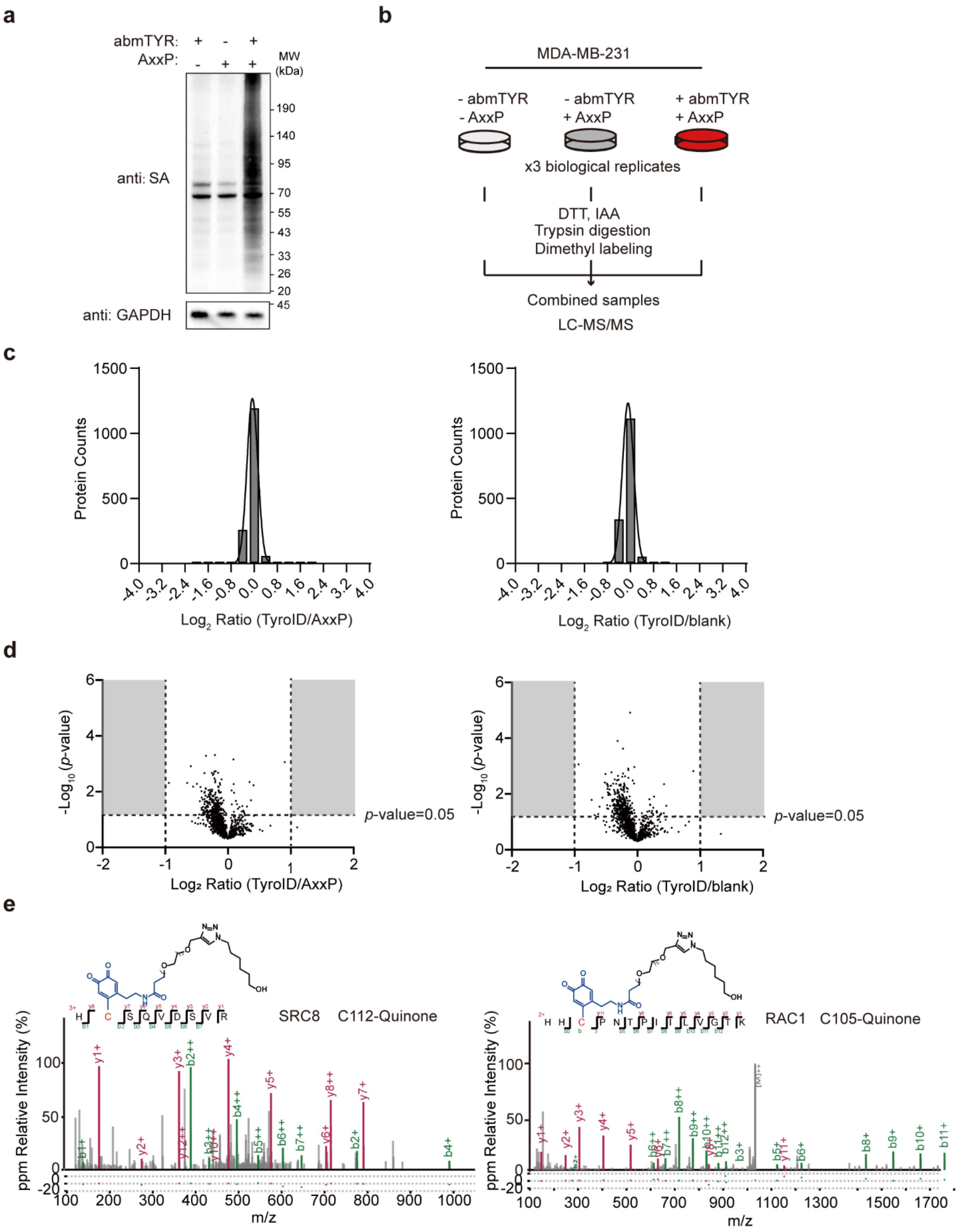


**Figure S5. Proteomic profiling of extracellular proteins by TyroID.** **a.** Validation of TyroID labeling by streptavidin blotting. **b.** The schematic of the global proteomic analysis to assess the impact of TyroID labeling on protein abundance. The total cell lysates from TyroID-labeled cells and control samples were directly subjected to a dimethyl labeling-based global proteomic analysis. **c.** The distribution of dimethyl labeling ratios (left: TyroID versus AxxP; right: TyroID versus blank). **d.** Volcano plots showing the dimethyl labeling ratios and *p*-values. Areas with a fold-change over 2 and *p*-value below 0.05 are gray. **e.** Tandem MS spectra of abmTYR-labeled peptides from cell-surface proteins with quinone modifications.

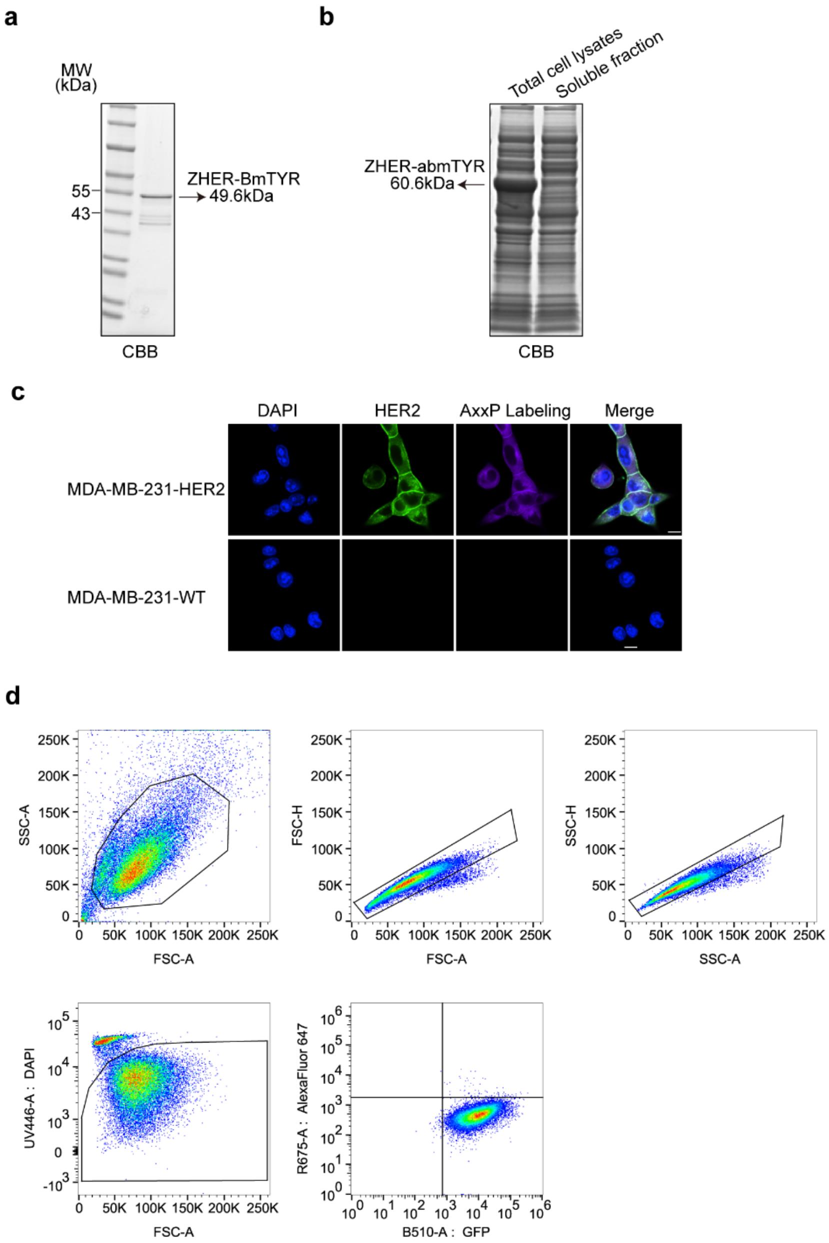


**Figure S6. Cell-selective extracellular protein labeling via HER2-targeting TyroID. a.** Purification of BmTYR fused with a HER2 nanobody (ZHER-BmTYR). **b.** Expression of the H subunit from abmTYR, fused with ZHER. **c.** Confocal fluorescence imaging of ZHER-BmTYR labeling on HER2-positive and HER2-negative MDA cells. HER2-positive and HER2-negative MDA cells were treated with 75 nM abmTYR and 50 μM AxxP for 30 minutes. The cells were fixed and click with azide-biotin after cell fixation to visualize the labeled proteins with streptavidin-AF647. Scale bars, 10 μm. **d.** Example gating strategy for flow cytometry analysis of MDA-MB-231 cells labeled by ZHER-BmTYR and BP. MDA-MB-231 cells were gated by Side scatter (SSC), Forward scatter (FSC) to select for the right single cell population and gated by DAPI to select for the living cells.

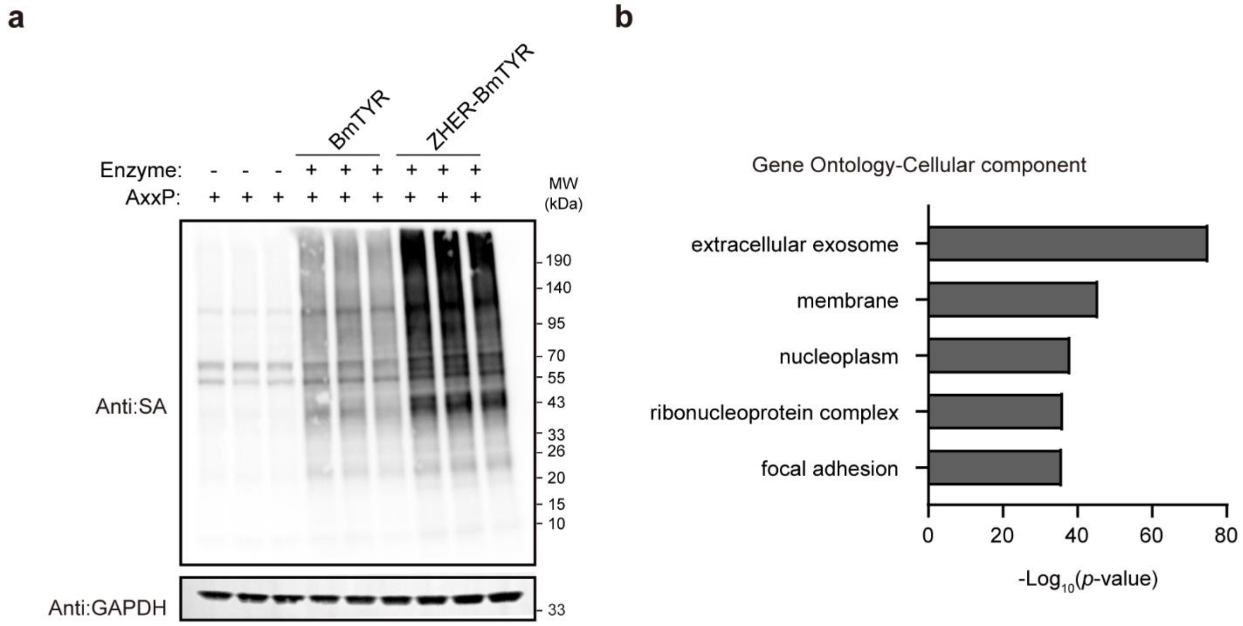


**Figure S7. Proteomic profiling of HER2-neighboring proteins by TyroID. a.** Validation of TyroID labeling on HER2-positive cells by streptavidin blotting. **b.** GO cellular component analysis of TyroID-enriched proteins.

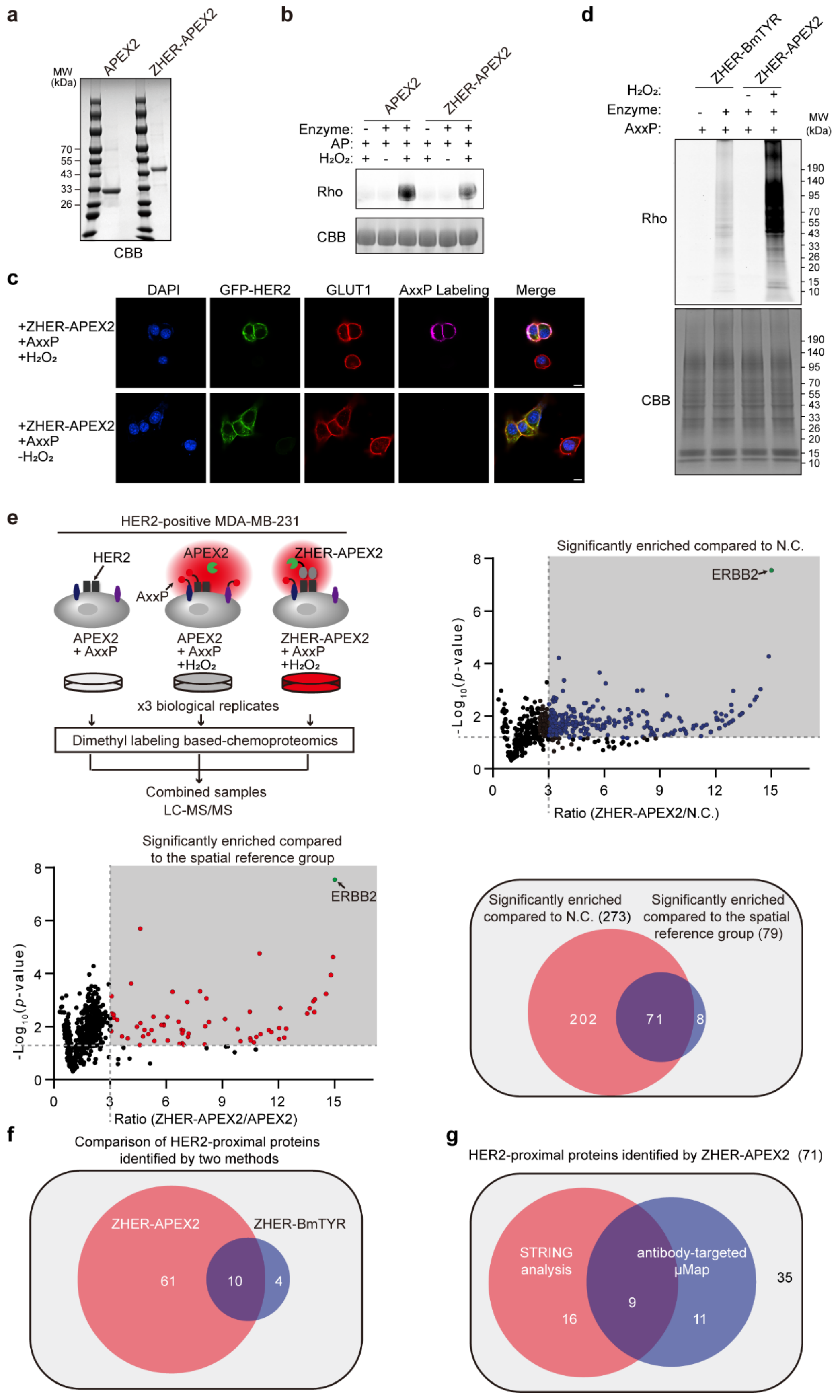
a
APEX2
ZHER-APEX2
MW (kDa)
70
55
43
33
26
CBB
b
APEX2
ZHER-APEX2
Enzyme:
AP:
H₂O₂:
Rho
CBB
c
DAPI
GFP-HER2
GLUT1
AxxP Labeling
Merge
+ZHER-APEX2 +AxxP +H₂O₂
+ZHER-APEX2 +AxxP -H₂O₂
d
ZHER-BmTYR
ZHER-APEX2
H₂O₂:
Enzyme:
AxxP:
MW (kDa)
Rho
CBB
190
140
95
70
55
43
33
26
20
15
10
e
HER2-positive MDA-MB-231
HER2
APEX2
ZHER-APEX2
AxxP
APEX2 + AxxP
APEX2 + AxxP +H₂O₂
ZHER-APEX2 + AxxP +H₂O₂
x3 biological replicates
Dimethyl labeling based-chemoproteomics
Combined samples
LC-MS/MS
Significantly enriched compared to N.C.
ERBB2
-Log₁₀(p-value)
Ratio (ZHER-APEX2/N.C.)
Significantly enriched compared to the spatial reference group
ERBB2
-Log₁₀(p-value)
Ratio (ZHER-APEX2/APEX2)
Significantly enriched compared to N.C. (273)
Significantly enriched compared to the spatial reference group (79)
202
71
8
f
Comparison of HER2-proximal proteins identified by two methods
ZHER-APEX2
ZHER-BmTYR
61
10
4
g
HER2-proximal proteins identified by ZHER-APEX2 (71)
STRING analysis
antibody-targeted μMap
35
9
16
11

**Figure S8. Proteomic profiling of HER2-neighboring proteins by ZHER-APEX2. a.** Purification of APEX2 and ZHER-APEX2 (APEX2 fused with the HER2 nanobody ZHER). **b.** The $H_2O_2$-dependent labeling of bovine serum albumin (BSA) by ZHER-APEX2 and AP *in vitro*. Recombinant BSA was treated with 75 nM APEX2/ZHER-APEX2 and 50 μM AP for 30 minutes, followed by 1 mM $H_2O_2$ for 1 minute, and then subjected to click reaction with azide-Rhodamine. The labeling intensity was determined by in-gel fluorescence scanning and Coomassie brilliant blue (CBB) staining was used to demonstrate the equal loading. **c.** Confocal fluorescence imaging of APEX2 labeling in a co-culture system. HER2-positive and HER2-negative MDA cells were mixed at a 1:20 ratio, followed by the treatment of 75 nM ZHER-APEX2 and 50 μM AxxP for 30 minutes, followed by 1 mM $H_2O_2$ for 1 minute. The cells were clicked with azide-biotin and then fixed with 4% PFA to visualize the labeled proteins with streptavidin-AF647. Scale bars, 10 μm. **d.** Comparison of in-gel fluorescence between ZHER-APEX2 labeling and ZHER-BmTYR labeling. **e.** Dimethyl labeling-based quantitative proteomics to map HER2-proximal proteins using ZHER-APEX2. The labeling group was compared against the negative control and spatial reference, respectively. Proteins enriched in both comparisons were assigned as HER2-proximal proteins. **f.** Overlap of HER2-proximal proteins identified by ZHER-BmTYR and ZHER-APEX2 labeling. **g.** Comparison of HER2-neighboring proteins identified by ZHER-APEX2 and μMAP.

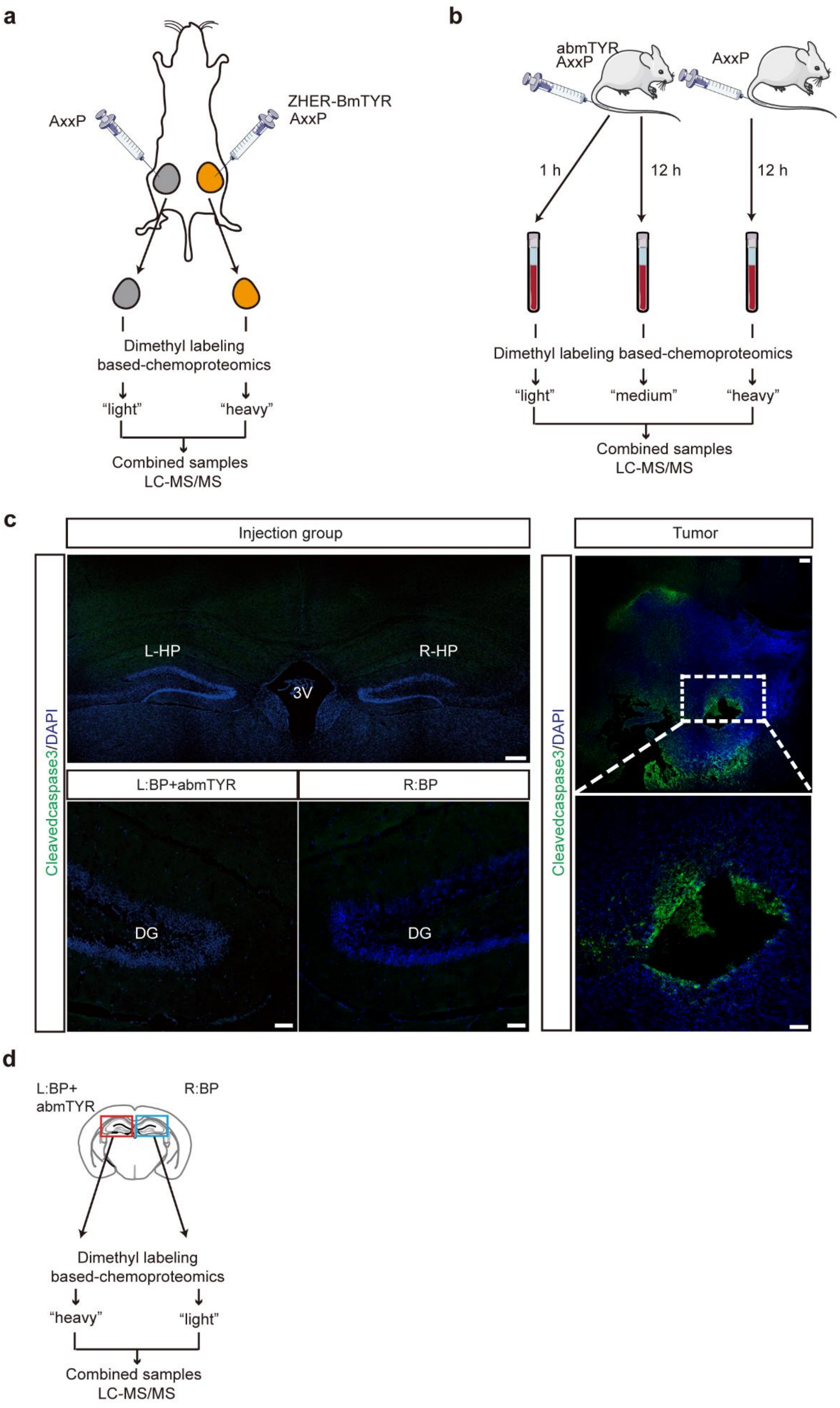
a
AxxP
ZHER-BmTYR
AxxP
Dimethyl labeling
based-chemoproteomics
"light"
"heavy"
Combined samples
LC-MS/MS
b
abmTYR
AxxP
AxxP
1 h
12 h
12 h
Dimethyl labeling based-chemoproteomics
"light"
"medium"
"heavy"
Combined samples
LC-MS/MS
c
Injection group
Tumor
Cleavedcaspase3/DAPI
L-HP
R-HP
3V
L:BP+abmTYR
R:BP
DG
DG
Cleavedcaspase3/DAPI
d
L:BP+
abmTYR
R:BP
Dimethyl labeling
based-chemoproteomics
"heavy"
"light"
Combined samples
LC-MS/MS

**Figure S9. *In vivo* labeling applications enabled by TyroID. a-b.** The workflow of dimethyl labeling-based quantitative proteomics for mapping HER2-neighboring proteins (**a**), tracking the turnover of plasma proteins (**b**). **c.** Histological staining of cleaved caspase 3 in the hippocampus (HP) and dentate gyrus (DG) of brains. The left (L) HP was injected with abmTYR with biotin phenol (BP), whereas the right (R) HP was injected with BP alone. The brain transplanted with GL261 glioma cells was used as a positive control for cleaved caspase 3 staining. **d.** The workflow of dimethyl labeling-based quantitative proteomics for identifying hippocampus-specific proteins *in vivo* by TyroID.

### Supplementary Tables

**Table S1. BmTYR-labeled and APEX2-labeled sites identified by superTOP-ABPP. Related to Figure 2.** Peptides modified by mito-BmTYR are shown in Tab1 with both phenol and quinone modifications on cysteines, histidines and lysines. Peptides with tyrosines modified by mito-APEX2 are shown in Tab 2. The modified residues are shown in lowercase letters.

**Table S2. Extracellular proteins labeled by abmTYR with AxxP. Related to Figure 4.** The TyroID/AxxP and TyroID/abmTYR ratios for each abmTYR-labeled protein are shown. The mean values of ratios and *p*-values were calculated based on three replicates.

**Table S3. Whole-cell proteomics quantifying protein abundance changes upon TyroID labeling.** Proteins quantified in all three biological replicates are shown with dimethyl labeling ratios (TyroID/AxxP and TyroID/blank) and *p*-values.

**Table S4. abmTYR-labeled sites identified by superTOP-ABPP. Related to Figure 4.** Peptides modified by abmTYR with AxxP are shown with both phenol and quinone modifications on cysteines, histidines and lysines. The modified residues are shown in lowercase letters.

**Table S5. HER2-proximal proteins labeled by ZHER-bmTYR and ZHER-APEX2 with AxxP in cell culture. Related to Figure 6 and S8.** The TyroID/N.C. and TyroID/spatial reference ratios for each labeled protein are shown. The TyroID-labeled sample was firstly compared to the negative control and proteins with TyroID/N.C. ratio over 3 and *p*-values below 0.05 were assigned as extracellular proteins, which are listed in Tab 1. The TyroID-labeled sample was then compared to the untargeted bmTYR ("spatial reference") and extracellular proteins with TyroID/spatial reference ratio over 3 and *p*-values below 0.05 were assigned as HER2-proximal proteins, which are listed in Tab 2. ZHER-APEX2 labeling samples were analyzed in the same way, with extracellular proteins and HER2-proximal proteins listed in Tab 3 and Tab 4, respectively.

**Table S6. HER2-proximal proteins labeled by ZHER-bmTYR with AxxP in tumor xenografts. Related to Figure 7.** The heavy/light ratio for each labeled protein is shown. The mean value and standard deviation were calculated based on three replicates.

**Table S7. Plasma proteins labeled by abmTYR with AxxP in the blood of living mice. Related to Figure 7.** The light/heavy ratio and medium/light for each labeled protein are shown. The mean value and *p*-value were calculated based on three replicates. The 1-hour labeling sample was firstly compared to the negative control and proteins with light/heavy ratio over 1.5 were assigned as enriched proteins. Then the 12-hour labeling sample was compared to the 1-hour labeling sample to quantify the turnover rates of labeled proteins.

**Table S8. Hippocampus-specific proteins labeled by abmTYR with BP. Related to Figure 8.** The heavy/light ratio for each labeled protein is shown. The mean value and standard deviation were calculated based on three replicates.
